# SPSID: A single-parameter shrinkage inverse-diffusion for denoising gene-regulatory networks

**DOI:** 10.64898/2026.01.19.700249

**Authors:** Hao Chen, Ge Han, Wenze Ding, Clara Grazian

## Abstract

Inferring gene regulatory networks (GRNs) from expression data is a fundamental problem in systems biology, but its accuracy is often undermined by structural noise arising from transitive correlations. These indirect interactions can obscure the true regulatory architecture, leading to a high rate of false positives. To address this, we introduce SPSID (Single-Parameter Shrinkage Inverse-Diffusion), a novel and robust network denoising framework. SPSID employs a principled spectral filter, built upon a shrinkage-regularized inverse-diffusion operator, to mathematically distinguish direct, one-step interactions from multi-step, indirect paths. This approach guarantees numerical stability and, through a fixed default parameter, effectively eliminates the need for data-dependent tuning. We conducted a comprehensive evaluation of SPSID on both extensive simulations and the gold-standard DREAM5 benchmark. The results demonstrate that SPSID outperforms state-of-the-art baseline methods in both AUROC and AUPR, exhibiting good stability across diverse network conditions. Furthermore, it functions as a post-processing tool, elevating the performance of multiple upstream GRN inference methods. By providing a computationally efficient and parameter-free solution to filter structural noise, SPSID offers a readily applicable tool for uncovering the underlying topology of complex biological networks with greater fidelity.

## I. Introduction

Transitive correlation is a fundamental concept in statistics, referring to the phenomenon where the association between two variables is mediated by one or more intermediate variables. In high-dimensional biological data, such as gene expression profiles [1], protein residue interactions [2], [3], [4], and other molecular features [5], transitive effects are ubiquitous and can lead to indirect relationships that obscure the underlying direct causal connections [6]. Without careful treatment, these indirect associations may result in misleading biological interpretations.

Besides classical measurement noise arising from sequencing depth, batch effects and other technical factors [7], Gene regulatory networks (GRNs) inference suffers from a second, often dominant, source of error, structural noise. Structural noise manifests as spurious edges created by the propagation of correlation along multi-step paths in the gene–gene interaction graph. A series of diffusion-based filters, including Network Deconvolution [8], Network Enhancement [9], and the recent RENDOR framework [10], were explicitly designed to attenuate these indirect links. Gene regulatory networks (GRNs) exemplify the complexity of biological systems, as a relatively small set of transcription factors (TFs) regulate the expression of numerous target genes by binding to specific DNA regulatory elements [11], [12]. Deciphering GRNs is crucial for understanding cellular function, elucidating the mechanisms underlying functional specialization, and uncovering the molecular basis of diseases[13], [14], [15], [16]. However, inferring GRNs is challenging due to the high dimensionality of the data and the prevalence of transitive correlations.

Recent advances in biotechnology and artificial intelligence have spurred the development of computational approaches for GRN inference from multi-omics data [17], [18], [19]. Many existing methods rely on pairwise correlation or mutual information as surrogates for causation, but these measures are prone to indirect associations. For instance, if gene A is correlated with gene B and gene B with gene C, the observed correlation between gene A and gene C might be an indirect effect mediated by gene B[20], [21]. Misinterpreting these transitive effects can lead to an elevated number of false positive regulatory interactions.

Here, we introduce SPSID (Single-Parameter Shrinkage Inverse-Diffusion), a novel network-denoising method that uses a shrinkage inverse diffusion operator to distinguish direct interactions from spurious, multi-step correlations. Unlike complex models requiring extensive tuning, SPSID is a powerful spectral filter built on a single-parameter regularization framework. We demonstrate its superiority and robustness through a two-pronged evaluation. In extensive simulations, SPSID consistently outperforms state-of-the-art baselines in area under the receiver-operating characteristic (AUROC) and area under the precision-recall curve (AUPR) with remarkable stability. Critically, on the gold-standard DREAM5 (Dialogue for Reverse Engineering Assessments and Methods 5) benchmark [22], it not only achieves top-ranked performance but also acts as a post-processing tool, significantly boosting the accuracy of multiple upstream GRN inference methods.

In the following sections, we detail the proposed SPSID algorithm, present a comprehensive performance evaluation on both simulated and real-world benchmark data, and discuss the implications of our findings for network inference. The remainder of this paper is organized as follows. Section IV presents an extensive simulation study, evaluating SPSID’s performance under various network conditions and comparing it against baseline methods. In Section V, we further validate our approach using the gold-standard DREAM5 benchmark datasets. Finally, we discuss our findings in Section VI and provide concluding remarks in Section VII.

## II. Related Work

The inference of gene regulatory networks is a long-standing and active area of research in computational biology. Over the years, a diverse array of methods has been developed to tackle this challenge, which can be broadly categorized based on their underlying principles.

### A. Gene Regulatory Network Inference Methods

Early and influential approaches relied on information theory, such as ARACNE, which uses the Data Processing Inequality to eliminate indirect interactions inferred from mutual information scores [23]. This line of MI-based research has continued to evolve, for instance, through the development of consensus-based algorithms and improved conditional mutual information tests [24]. Another major category consists of regression-based methods. These approaches frame the problem as a feature selection task, where the expression of each gene is modeled as a function of all potential regulators (TFs). Prominent examples include TIGRESS [25] and tree-based ensemble methods like GENIE3 [26] and GRNBoost2 [27], which have demonstrated robust performance in various benchmarks. More recently, with the rise of deep learning, methods employing convolutional neural networks [28], [29], [30] and graph neural networks [31], [32] have been proposed to capture complex, non-linear dependencies in gene expression data. While these methods vary in their approach, they all produce a weighted adjacency matrix or a ranked list of potential regulatory interactions. This output, however, is invariably affected by statistical noise and, more critically, structural artifacts inherent in the data.

### B. The Challenge of Transitive Correlations

A primary source of error in GRN inference is the prevalence of transitive correlations, also known as indirect effects [33], [21], [34]. If a transcription factor A regulates gene B, and B in turn regulates gene C, a strong statistical association may be observed between A and C, even in the absence of a direct causal link. This phenomenon can lead to a proliferation of false-positive edges, obscuring the true network topology. Statistically, this issue is related to the problem of distinguishing marginal from conditional dependence. Gaussian Graphical Models (GGMs), for instance, use partial correlations to infer conditional independence and thus identify direct links in a network, with methods like glasso providing efficient algorithms for sparse inverse covariance estimation [35], [36], [37]. However, GGMs rely on assumptions of normality and linearity which may not hold for complex biological data [38], [39], [40]. The challenge, therefore, is to develop methods that can effectively disentangle these direct and indirect effects in a more general, non-parametric setting.

### C. Network Denoising and Deconvolution

To specifically address the problem of structural noise, a class of post-processing methods focused on network denoising and deconvolution has emerged. These methods take a raw, noisy interaction matrix as input and apply a filter to refine the scores. A seminal idea in this space is Network Deconvolution (ND), which models the observed network as the result of a matrix power series (a transitive closure process) over the true direct-interaction network and seeks to invert this process [8]. Another approach, proposed by Barzel and Barabási, uses a “global silencing” operator (Silencer) to suppress indirect correlations based on the network’s global structure [20].More recent work has leveraged concepts from graph theory and diffusion processes. Network Enhancement (NE) uses an iterative diffusion process on a doubly stochastic matrix to sharpen the network structure [9]. The idea of information diffusion on graphs has a rich history, with methods like diffusion kernels [41], [42], [43] and Random Walk with Restart (RWR) [44], [45], [46] being used for tasks like node ranking and link prediction. The recently proposed Reverse Network Diffusion On Random Walks (RENDOR) method builds on this, using a geometric series-based diffusion model that is inverted to recover direct interactions [10]. While powerful, many of these methods either rely on specific modeling assumptions, require the tuning of sensitive, data-dependent hyperparameters, or lack guaranteed numerical stability. Our proposed method, SPSID, contributes to this line of work by introducing a shrinkage-regularized inverse operator. This formulation provides a mathematically robust and parameter-free framework to filter transitive effects, directly addressing the key limitations of prior approaches.

## III. The SPSID Denoising Methodology

Our proposed SPSID framework denoises a raw gene-regulatory score matrix, *W*_obs_, through the three-stage process illustrated in Fig. 1. First, we perform a series of pre-processing steps to transform the raw scores into a mathematically well-posed, row-stochastic transition matrix, *P*_obs_. Second, we apply the core shrinkage-regularised inverse-diffusion (SPSID) operator to *P*_obs_ to filter out indirect, transitive interactions and yield a denoised matrix, *P*_dir_. Finally, we conduct a node-specific re-weighting using the network’s stationary distribution to restore biological context and produce the final output, *W*_dir_. The details of each stage are described below.

**Fig. 1:**
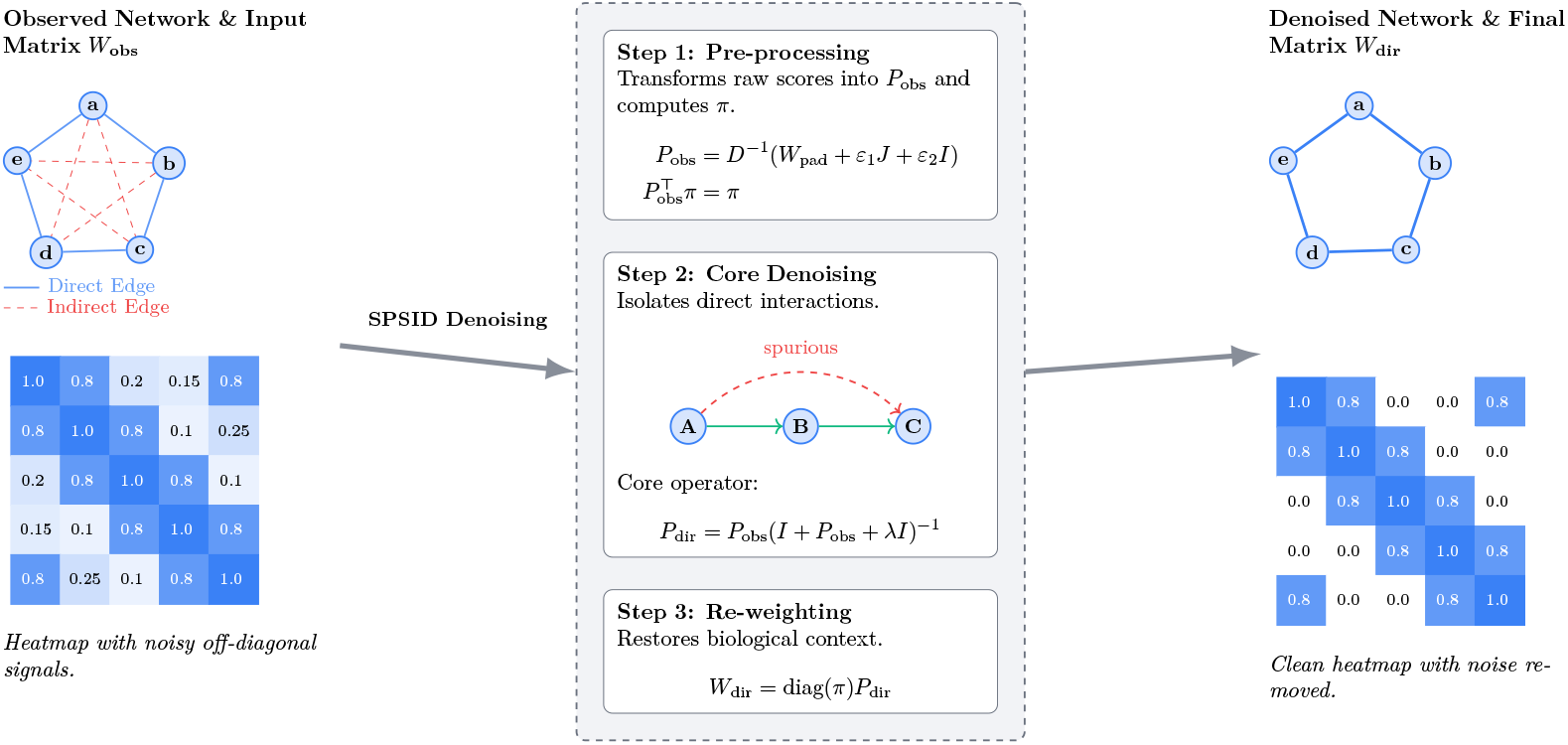
An overview of the three-stage SPSID denoising methodology. The process begins with a raw, noisy observed network (left), which is transformed by the central framework to isolate direct interactions and produce a clean, re-weighted final network (right).

### A. Step 1: Pre-processing for a Well-Posed Transition Matrix

Let *W*_obs_ ∈ ℝ^*m*×*n*^ be the weighted TF–gene score matrix obtained from an upstream GRN-inference engine (e.g. GE-NIE3, GRNBoost2), where *m* is the number of transcription factors (rows) and *n* is the total number of genes (columns). The column set therefore contains the same *m* TFs plus the *n* − *m pure target* genes that never act as regulators. Because the downstream steps require square matrices, we first embed *W*_obs_ into an *n* × *n* square matrix.

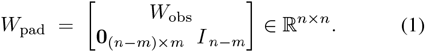

This padding operation transforms the original *m*×*n* matrix into a square *n* ×*n* matrix. The lower-right identity block *I*_*n*−*m*_ assigns a self-loop of 1 to each pure target, guaranteeing at least one non-zero entry per row and preventing division-by-zero during subsequent normalisation.

Even after padding, *W*_pad_ can be sparse and may contain zero rows or bipartite structures, which lead to ill-conditioned operators [10]. We stabilise the matrix by

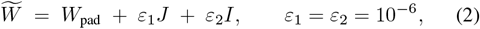

where *J* and *I* denote the all-ones matrix and the identity matrix, respectively. The dual *ε*-perturbation, composed of a global lift (*ε*_1_*J*) and a local lift (*ε*_2_*I*), ensures that the subsequent random-walk matrix is irreducible and aperiodic. These properties are purely for numerical stability and do not alter the biological ranking of edge weights, as the perturbation constants are negligible.

To obtain a probabilistic view of information flow, we normalise the stabilised weight matrix row-wise. Let

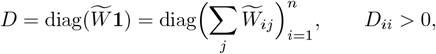

and define the transition matrix

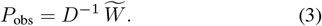

Every row of *P*_obs_ sums to one, so *P*_obs_(*i, j*) is the probability of a one-step move from source gene *i* to target *j*. Because *P*_obs_ is irreducible and aperiodic, the random walk it describes possesses a unique stationary distribution. This is a probability vector *π* representing the long-run behaviour of the network, where each element *π*_*i*_ is the proportion of time a random walker will spend at gene *i* after many steps. Concretely, we define *π* as the column vector

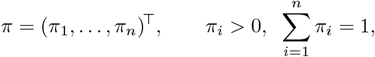

which uniquely satisfies the equilibrium equation:

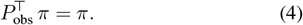

Because *P*_obs_ is irreducible and aperiodic, it admits a unique stationary distribution *π* satisfying 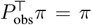 and **1**^*T*^*π* = 1 [47]. We obtain *π* with the standard power-iteration method [48]. This pre-processing stage transforms an arbitrary weight matrix into a well-conditioned Markov kernel, laying the mathematical foundation for the next step.

### B. Step 2: The Core SPSID Operator for Denoising

The row-stochastic matrix *P*_obs_ encodes both direct (1-step) and indirect (≥2-step) gene influences. An indirect influence arises from transitive effects within the network; for instance, if a transcription factor A directly regulates gene B, and gene B in turn directly regulates gene C (a path represented as A → B → C), a strong statistical correlation may be observed between A and C, even in the absence of a direct causal link. These transitive effects are a major source of spurious edges in inferred networks. In this context, the “noise” we aim to filter is precisely this web of indirect influences that obscures the true underlying network structure. The core challenge, therefore, is to mathematically deconvolve the observed matrix *P*_obs_ to isolate the matrix of true direct interactions, *P*_dir_.

This conceptual decomposition can be expressed as:

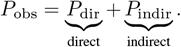

A prominent classical approach is Neumann-series deconvolution [8]. This framework models the observed network *P*_obs_ as arising from the true direct network *P*_dir_ via a transitive closure process,

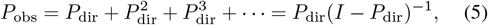

leading to the closed-form inverse solution:

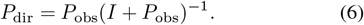

This model posits that the observed interactions are the sum of effects propagated along all paths of length one or more in the true direct-interaction network, *P*_dir_. The core task of denoising, therefore, is to invert this formula to solve for the underlying *P*_dir_ given the observed matrix *P*_obs_. However, the validity of this method hinges on a critical pre-processing step: the input matrix *P*_obs_ must be linearly scaled to ensure that the eigenvalues of the inferred direct matrix lie within the convergence interval (−1, 1). This scaling factor is a data-dependent hyperparameter, and the eigenvalue-based approach is most directly applicable to symmetric matrices.

A more recent paradigm is to model the observed network as a result of a geometrically damped diffusion process. For instance, the RENDOR method [10] proposes a forward model that leads to the following inverse diffusion operator for denoising. The operator *f*_*m*_(*P*) arises from a normalized geometric expansion of higher-order transitions, where the contribution of each path of length *k* is weighted by 1*/m*^*k*^. The closed-form expression is derived by recognizing this expansion as a convergent matrix power series:

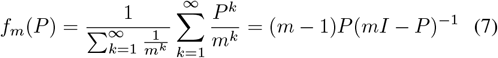

This guarantees spectral stability for all *m>* 1, and allows interpretation of *m* as a damping factor controlling the trade-off between direct and transitive effects.

The RENDOR [10] diffusion operator *f*_*m*_(*P*) is originally defined on the true direct-transition matrix *P*_dir_, modeling the forward process that generates the observed matrix *P*_obs_:

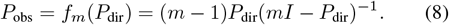

In practice, however, we only observe *P*_obs_, and aim to recover *P*_dir_. Inverting the forward diffusion process yields the denoising operator:

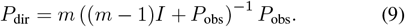

Here, *m >* 1 controls the suppression of higher-order transitive effects. While effective, this approach relies on a hyperparameter *m >* 1 to control the damping, which must be chosen empirically. Furthermore, this operator does not guarantee the non-negativity of the output matrix, requiring heuristic post-processing steps to correct negative entries.

The observed matrix *P*_obs_ mixes direct, one–step influences with indirect, multi–step (transitive) influences that manifest as statistical associations. As established by the Neumann–series/deconvolution identities presented just above, denoising can be viewed as an inverse problem: recover the one–step component while attenuating contributions from paths of length ≥ 2. While classical approaches like Neumann-series deconvolution and other diffusion models exist [8], [10], they often require data-dependent hyperparameter tuning or lack guaranteed numerical stability. The operator introduced below (Eq. (10)) implements precisely this inverse view but places a single shrinkage term inside the inverse. This has two immediate benefits without any data–dependent tuning: (i) it guarantees numerical well–conditioning of the inversion and (ii) it preferentially suppresses higher–order (indirect) modes relative to the one–step (direct) mode, thereby separating true transitive effects from mere statistical associations. For numerical hygiene, we clip negligible negative entries (from floating–point round–off) before the final re–weighting step. In this work, we propose SPSID, a novel inverse diffusion operator that incorporates a Tikhonov shrinkage regulariser term directly into the operator:

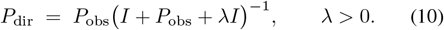

The shrinkage term *λI* provides two complementary benefits. First, it stabilises the inversion.

Let 𝒜 = (1+*λ*)*I* +*P*_obs_ be the matrix to be inverted. We can rewrite this as 𝒜 = (1 + *λ*)(*I* + *Q*), where *Q* = *P*_obs_*/*(1 + *λ*).

Since *P*_obs_ is row-stochastic, its *L*_∞_-norm (maximum absolute row sum) is ||*P*_obs_||_∞_ = 1. This implies ||*Q*||_∞_ = ||*P*_obs_ ||_∞_*/*(1+*λ*) = 1*/*(1+*λ*). As *λ >* 0, we have ||*Q*||_∞_ *<* 1, which guarantees the convergence of the Neumann series for (*I* + *Q*)^−1^.

We can now bound the norm of the inverse, ||𝒜^−1^||_∞_:

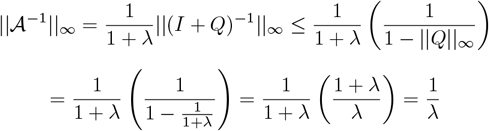

The norm of the original matrix is bounded by ||𝒜||_∞_ ≤ ||(1 + *λ*)*I*||_∞_ + ||*P*_obs_||_∞_ = (*λ* + 1)+1 = *λ* + 2.

Finally, the condition number is bounded by:

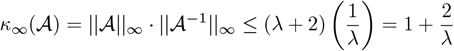

With our fixed default of *λ* = 10^3^, the condition number *κ*_∞_(𝒜) ≤ 1.002. This is extremely close to 1, guaranteeing that the matrix is exceptionally well-conditioned and the inverse is numerically stable.

Second, *λI* introduces a mode-dependent shrinkage that separates direct from indirect influence. Under the SPSID transform, an eigenvalue *μ* of *P*_obs_ is scaled by the function *g*(*μ*) = *μ/*(1 + *μ* + *λ*). The eigenvalue *μ* = 1, which corresponds to direct interactions, is scaled by *g*(1) = 1*/*(2 +*λ*). In contrast, any indirect mode with |*μ*| *<* 1 is suppressed much more significantly relative to the direct mode. The ratio of suppression is

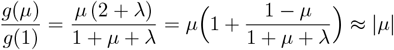

for large *λ*.

With our fixed default of *λ* = 10^3^, indirect components are sharply attenuated while the internal ranking of direct interactions is preserved. This eliminates the need for hyper-parameter tuning. The inverse in Equation (10) is computed efficiently via Lower–Upper (LU) factorisation, with a negligible runtime of *O*(*n*^3^) for typical network sizes. In addition, in our implementation, any numerically negligible negative entries (≈ −10^−12^) arising from floating-point imprecision in *P*_dir_ are clipped to zero before the final re-weighting step.

### C. Step 3: Post-processing and Final Score Re-weighting

The diffusion step produces *P*_dir_, a row–sub-stochastic *square* matrix (*n* × *n*). To restore biologically meaningful magnitudes and incorporate global node importance, we modulate each row by the long-run influence of its source gene, captured by the stationary distribution *π* from Step 1. We define the final denoised matrix as

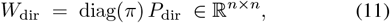

so that row *i* is multiplied by the scalar *π*_*i*_. Each transcription factor now retains a total *regulatory budget* proportional to its steady-state mass *π*_*i*_.

Biologically, this re-weighting serves three purposes: it integrates global importance, avoids degree bias, and maintains comparability of edge rankings. When the original input is rectangular (*m < n*), we simply extract the leading *m* × *n* block from *W*_dir_ to get the final TF–target matrix, ready for downstream analysis. The complete SPSID workflow is consolidated in Algorithm 1.

## IV. Simulation Results

### A. Simulation design

We consider networks of size *n* = 100 consisting of *m*_TF_ = 50 transcription factors (TFs) and *n* − *m*_TF_ = 50 pure target genes. For a prescribed sparsity level *ρ* each potential TF–target edge is drawn independently with probability *ρ* and assigned a weight *w*_*ij*_ ∼ Unif[0.5, 1]; self-loops are disallowed. The resulting weight matrix is denoted **G**_dir_. We vary the density over *ρ* ∈ *{*0.01, 0.03, 0.05, 0.07, 0.09} to emulate transcriptional programmes of different compactness.

#### Algorithm 1

SPSID: shrinkage inverse-diffusion for GRN denoising.

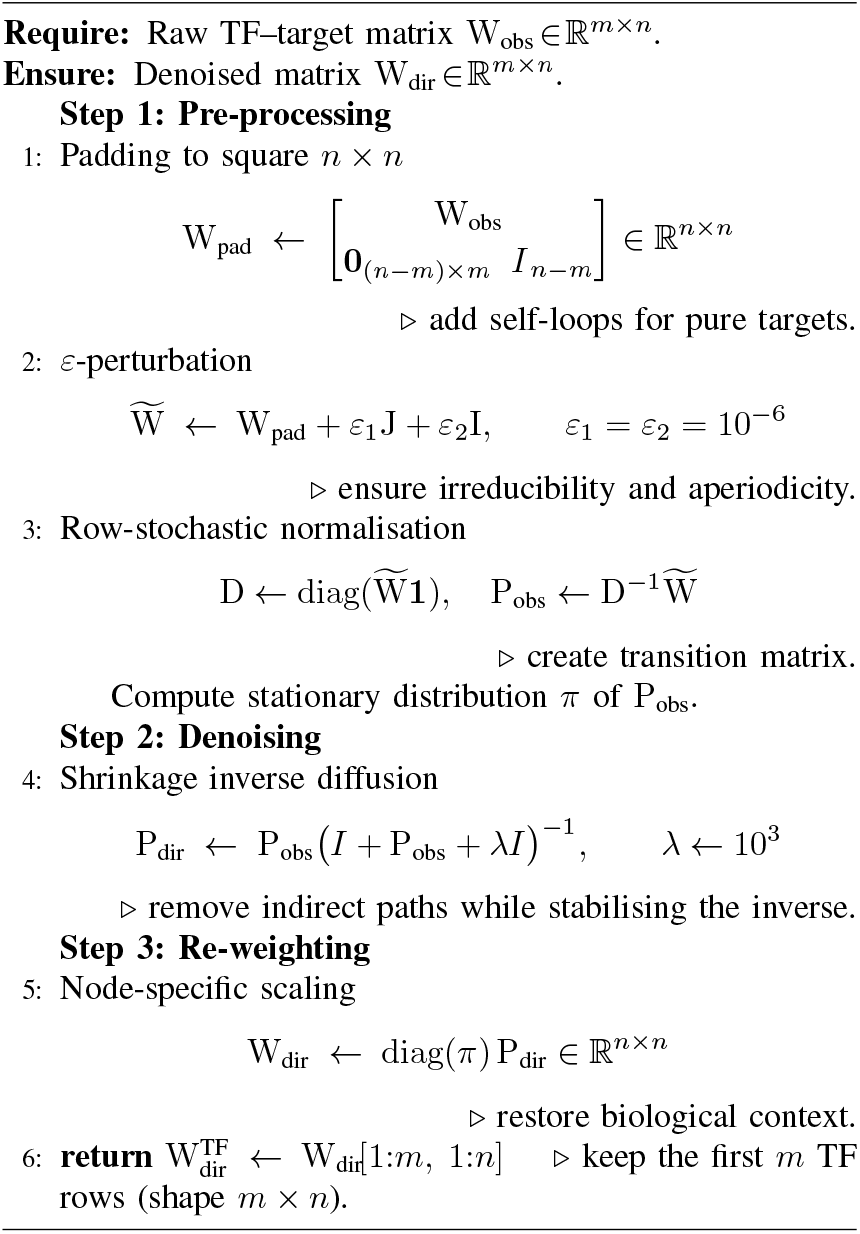

To mimic transitive correlations and experimental noise we generate an *observed* network

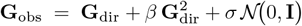

where (i) the quadratic term aggregates indirect two-step influences with strength parameter *β*; (ii) 𝒩 (0, **I**) is an i.i.d. Gaussian matrix, and negative entries are clipped to zero. We explore five noise levels *σ* ∈ {0.1, 0.3, 0.5, 0.7, 0.9} and three transitive strengths *β* ∈ {0.1, 0.3, 0.5, 0.7, 0.9}, while keeping *ρ* = 0.05 fixed when varying *β* or *σ*. For each triple (*ρ, β, σ*) we simulate *N*_rep_ = 1000 independent replicates, yielding 5 × 3 × 4 × 1000 = 60,000 observed networks in total. SPSID is executed with the default *λ* = 10^3^ and *ε*_1_ = *ε*_2_ = 10^−6^. For each replicate we compute (i) the area under the receiver-operating characteristic (AUROC); (ii) the area under the precision–recall curve (AUPR); and (iii) the number of true-positive (TP) edges retained among the top-*k* predictions for *k* ∈ {100, 200,…, 1500} . Edge probabilities are read directly from the denoised weight matrices, whereas ground-truth labels are obtained by binarising **G**_dir_. All metrics are averaged over replicates, and 95% confidence intervals use the normal approximation with unbiased sample variance. To assess the robustness of SPSID itself we perform an additional one-factor experiment in which *λ* ∈ {100, 200, 300,…, 1000, 5000, 10000} while fixing *ρ* = 0.05, *β* = 0.5 and *σ* = 0.5.

Every simulated network is post-processed by SPSID and five published denoising methods, RENDOR [10], Network Deconvolution (ND) [8], Network Enhancement (NE) [9], Global Silencing of Indirect Correlations (Silencer) [20], and Inverse Correlation Matrix (ICM) [49], so that all six methods are evaluated head-to-head under identical simulation conditions. Subsequent subsections present the baseline comparison (IV-B); an *extended-scenario analysis* that systematically varies network sparsity, noise level, and transitive strength (IV-C); the edge-retention analysis (IV-D); and the *λ*-sensitivity study (IV-E).

### B. Baseline performance analysis

Table I reports the mean AUROC/AUPR and the associated 95 % confidence intervals for the six denoising methods under the default simulation setting *ρ* = 0.05, *β* = 0.1, *σ* = 0.5 (*N*_rep_ = 1000 replicates). SPSID attains the highest values on both metrics, followed by RENDOR and ND; NE, Silencer and ICM perform substantially worse.

**TABLE I:**
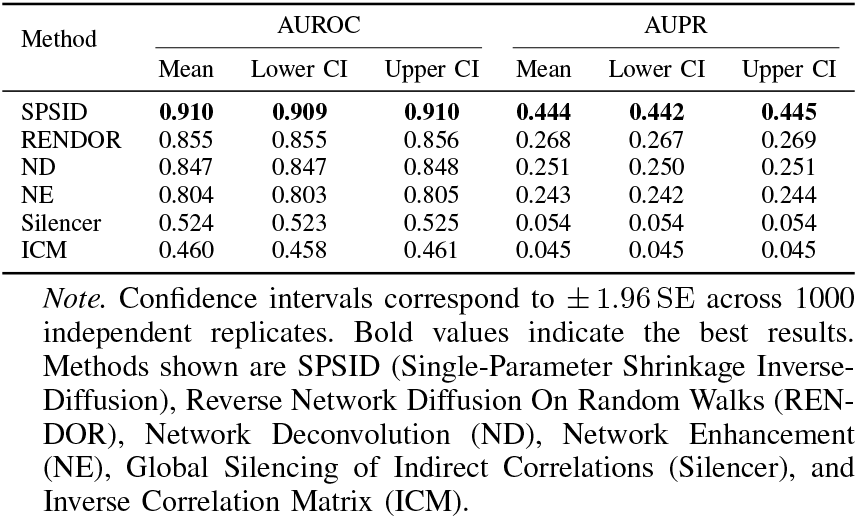
Performance comparison of network–denoising methods (*n*=100, *ρ*=0.05, *β*=0.1, *σ*=0.5)

| Method | AUROC |  |  | AUPR |  |  |
| --- | --- | --- | --- | --- | --- | --- |
|  | Mean | Lower CI | Upper CI | Mean | Lower CI | Upper CI |
| SPSID | **0.910** | **0.909** | **0.910** | **0.444** | **0.442** | **0.445** |
| RENDOR | 0.855 | 0.855 | 0.856 | 0.268 | 0.267 | 0.269 |
| ND | 0.847 | 0.847 | 0.848 | 0.251 | 0.250 | 0.251 |
| NE | 0.804 | 0.803 | 0.805 | 0.243 | 0.242 | 0.244 |
| Silencer | 0.524 | 0.523 | 0.525 | 0.054 | 0.054 | 0.054 |
| ICM | 0.460 | 0.458 | 0.461 | 0.045 | 0.045 | 0.045 |

*Note.* Confidence intervals correspond to $\pm 1.96$ SE across 1000 independent replicates. Bold values indicate the best results. Methods shown are SP SID (Single-Parameter Shrinkage Inverse-Diffusion), Reverse Network Diffusion On Random Walks (RENDOR), Network Deconvolution (ND), Network Enhancement (NE), Global Silencing of Indirect Correlations (Silencer), and Inverse Correlation Matrix (ICM).

Table I and Fig. 2 compare six denoising filters under the default simulation setting (*n*=100, *ρ* = 0.05, *β* = 0.1, *σ* = 0.5). SPSID attains the highest mean scores on both metrics (AUROC = 0.910, AUPR = 0.444), with 95 % confidence intervals that differ only in the third decimal place. Relative to the strongest baseline (RENDOR), SPSID improves AUROC by +6.4% and AUPR by an impressive +65.7 %. RENDOR (AUROC 0.855; AUPR 0.268) and ND (AUROC 0.847; AUPR 0.251) provide moderate gains over raw networks but fall well short of SPSID—in particular, their AUPR values lag by nearly 0.18. NE delivers only marginal benefit (AUROC 0.804; AUPR 0.243), while Silencer and ICM perform poorly (AUROC ≤ 0.524, AUPR ≤ 0.054), indicating that they not only fail to suppress indirect noise but may even degrade network quality. Overall, SPSID delivers the good accuracy and the consistency among all evaluated methods. The following subsections analyse robustness to network sparsity, noise level, and transitive strength (IV-C); the edge-retention analysis (IV-D); and the *λ*-sensitivity study (IV-E).

**Fig. 2:**
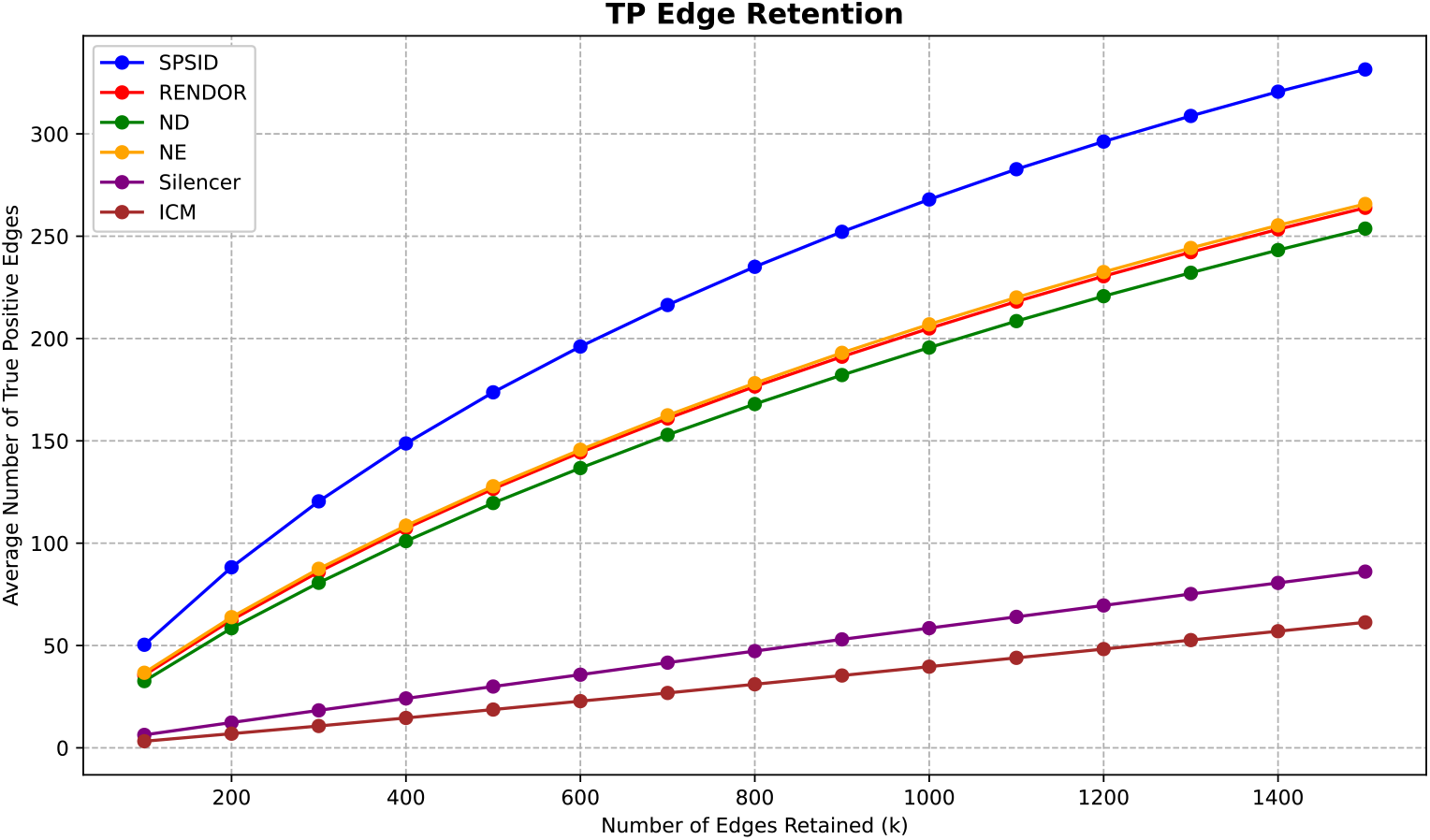
Performance comparison of denoising methods across 1000 simulation replicates. (A) Distribution of AUROC scores shown as a box plot. (B) Distribution of AUPR scores shown as a box plot. (C) Mean AUROC scores presented in a bar chart. (D) Mean AUPR scores presented in a bar chart. Methods shown are SPSID (Single-Parameter Shrinkage Inverse-Diffusion), Reverse Network Diffusion On Random Walks (RENDOR), Network Deconvolution (ND), Network Enhancement (NE), Global Silencing of Indirect Correlations (Silencer), and Inverse Correlation Matrix (ICM).

### C. Extended-scenario analysis

Beyond the default setting, we systematically investigate several factors. The main analysis varies three parameters (Network sparsity, Noise level, and transitive strength) while keeping others fixed. Specifically:

- **Network sparsity** — *ρ* ∈ { 0.01, 0.03, 0.05, 0.07, 0.09}, with *β* = 0.5 and *σ* = 0.5.
- **Noise level** — *σ* ∈ {0.1, 0.3, 0.5, 0.7, 0.9}, with *ρ* = 0.05 and *β* = 0.5.
- **Transitive strength** — *β* ∈ {0.1, 0.3, 0.5, 0.7, 0.9}, with *ρ* = 0.05 and *σ* = 0.5.

For every setting we simulate *N*_rep_ = 1000 networks and report the mean AUROC/AUPR. The results for these three main sweeps are in **Supplementary Table S2**. As *ρ* increases from 0.01 to 0.09 (**Supplementary Table S2a**), SPSID retains an AUROC above 0.83 and its AUPR almost doubles (0.177 → 0.317), consistently achieving the highest scores throughout the sweep. Increasing Gaussian noise (*σ* = 0.1 → 0.9) (**Supplementary Table S2b**) degrades every method, yet SPSID demonstrates strong robustness, preserving an AUROC of 0.758 and an AUPR of 0.156 even at the highest noise level. When quadratic influence (*β*_1_) grows from 0.1 to 0.9 (**Supplementary Table S2c**), all methods suffer in precision, but SPSID consistently maintains the top performance in both AUROC and AUPR across all levels of transitive noise.

To further probe the method’s robustness, we evaluated two additional challenging scenarios. First, we tested the extreme boundary case of the quadratic model at *β*_1_ = 1.0. The results **(Supplementary Table S3)** confirm the trend: SPSID remains the top performer, achieving the highest scores for both mean AUROC (0.828) and mean AUPR (0.172). Second, to test performance against more complex, higher-order noise, we introduced a new simulation model incorporating both quadratic and cubic terms, defined as:

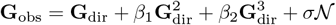

We fixed the quadratic noise (*β*_1_ = 0.5) and swept the new cubic noise parameter (*β*_2_). As shown in **Supplementary Table S4**, SPSID again outperforms all competitors across the entire sweep of cubic noise strengths. In summary, with a *single* hyper-parameter choice *λ*, SPSID delivers state-of-the-art performance across a broad spectrum of sparsity, noise, and transitive-effect levels, including boundary conditions and higher-order noise structures, mirroring practical GRN-inference scenarios where such factors are rarely known a priori.

### D. Edge Retention Analysis

Throughout this subsection a positive (or true) edge is a direct *TF* → *target* interaction that appears in the simulation ground-truth matrix; self-loops and target → TF links are excluded. A predicted edge is therefore counted as a true positive (TP) if the corresponding entry in **G**_dir_ equals 1. Table II and Fig. 3 report the *average* number of TPs recovered when only the top-*k* scores are retained (*k* ∈ {100, 300, 500, 700, 1000, 1500} ; *ρ* = 0.05, *β* = 0.5, *σ* = 0.5). For all six methods the mean TP count increases monotonically with *k*, yet the growth rate varies drastically. SPSID dominates at every threshold: at *k* = 100, it achieves 50 TPs (i.e., +39 % over RENDOR (36) and +52 % over ND (33)); at *k* = 500, it finds 174 TPs versus 128 (NE) and 127 (RENDOR); and at *k* = 1500, its 331 TPs exceed the runner-up RENDOR (264) by 67 and outscore all other baselines by at least 65. The consistently higher TP yield indicates that SPSID assigns larger scores to genuine direct interactions, thereby enriching the very top of the ranked list. This property is essential when downstream experimental validation can probe only a limited number of candidate edges.

**TABLE II:**
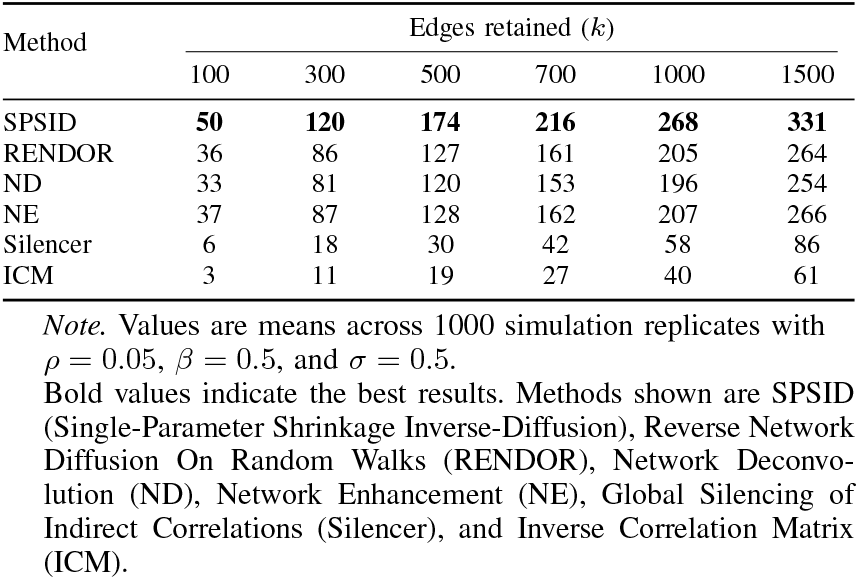
Average number of true–positive (TP) edges at selected retention thresholds.

**Fig. 3:**
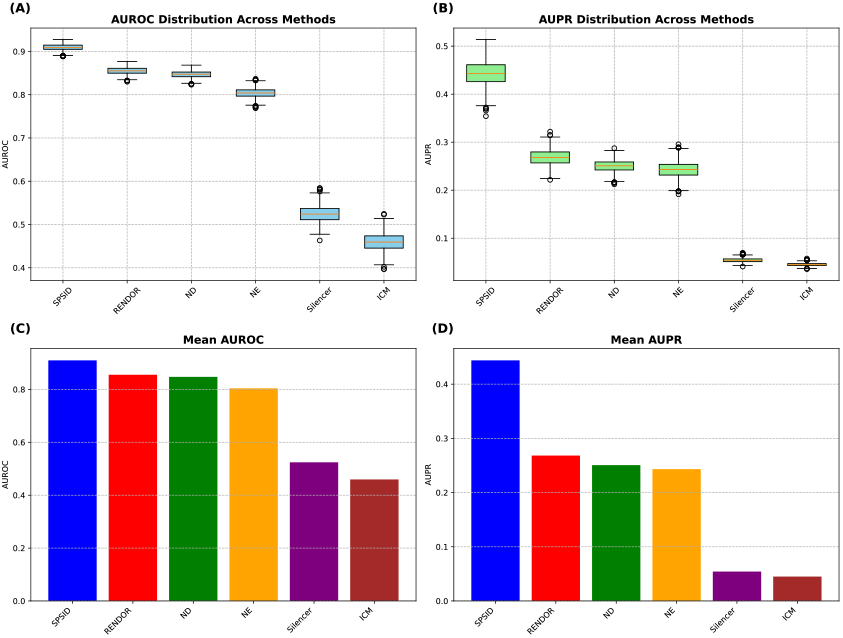
Comparison of the number of true positive (TP) edges when retaining different numbers of top-scoring edges (*k*) from 100 to 1500. Methods shown are SPSID (Single-Parameter Shrinkage Inverse-Diffusion), Reverse Network Diffusion On Random Walks (RENDOR), Network Deconvolution (ND), Network Enhancement (NE), Global Silencing of Indirect Correlations (Silencer), and Inverse Correlation Matrix (ICM).

### E. Sensitivity to the shrinkage parameter λ

To examine how strongly SPSID depends on its single hyper-parameter *λ*, we repeated the default experiment (*ρ* = 0.05, *β* = 0.5, *σ* = 0.5) for *λ* ∈ {100,…, 10000}, generating *N*_rep_ = 1000 replicates for each setting. Table III lists the resulting AUROC and AUPR together with their 95 % confidence intervals. AUROC values are consistently between 0.878 and 0.879, while AUPR values vary only between 0.304 and 0.305.These results confirm that SPSID is insensitive to the precise choice of *λ* provided it lies within a broad. We therefore keep the default *λ* = 10^3^ for all other experiments, ensuring maximal indirect-signal attenuation while preserving numerical stability.

**TABLE III:** SPSID sensitivity to shrinkage parameter *λ*.

| $\lambda$ | AUROC | | | AUPR | | |
| --- | --- | --- | --- | --- | --- | --- |
|  | Mean | Lower CI | Upper CI | Mean | Lower CI | Upper CI |
| 100 | 0.878 | 0.878 | 0.879 | 0.304 | 0.303 | 0.305 |
| 200 | 0.879 | 0.878 | 0.879 | 0.304 | 0.303 | 0.305 |
| 300 | 0.878 | 0.878 | 0.879 | 0.304 | 0.303 | 0.306 |
| 400 | 0.879 | 0.878 | 0.879 | 0.305 | 0.303 | 0.306 |
| 500 | 0.879 | 0.878 | 0.879 | 0.305 | 0.304 | 0.306 |
| 600 | 0.878 | 0.878 | 0.879 | 0.304 | 0.302 | 0.305 |
| 700 | 0.878 | 0.878 | 0.879 | 0.304 | 0.303 | 0.305 |
| 800 | 0.878 | 0.878 | 0.879 | 0.304 | 0.303 | 0.305 |
| 900 | 0.878 | 0.878 | 0.879 | 0.304 | 0.303 | 0.305 |
| 1000 | 0.879 | 0.878 | 0.879 | 0.304 | 0.303 | 0.306 |
| 5000 | 0.878 | 0.878 | 0.879 | 0.304 | 0.303 | 0.305 |
| 10000 | 0.878 | 0.878 | 0.879 | 0.304 | 0.303 | 0.306 |

Note. Confidence intervals are $\pm 1.96$ SE over 1000 replicates.

### F. Computational Performance and Scalability

Finally, we assessed the computational cost of SPSID. The *O*(*n*^3^) complexity, which is standard for methods involving matrix inversion, arises from the single inversion step in Equation 10. We benchmarked the wall-clock runtime of all denoising methods on synthetic networks of increasing size *n* (see Supplementary Table S1 for full results). The results in Table S1 show that SPSID’s absolute runtime remains practical for typical GRN sizes. On a standard desktop, SPSID takes on average 0.24 s (SD 0.02) at *n* = 500, 1.01 s (SD 0.02) at *n* = 1000, 5.33 s (SD 0.01) at *n* = 2000, 17.49 s (SD 0.17) at *n* = 3000, 48.47 s (SD 0.06) at *n* = 4000, and 148.72 s (SD 0.25) at *n* = 6000. SPSID is consistently faster than RENDOR at all sizes, and both are *orders of magnitude* faster than ND (e.g., ND requires 2373.05 s at *n* = 6000). Simpler baselines such as Silencer and ICM are the fastest in wall-clock time, which reflects their much cheaper update rules but also their lower accuracy. NE lies between SPSID/RENDOR and these low-cost baselines. Overall, SPSID offers competitive runtime among the higher-accuracy denoisers: its cubic cost matches that of other global diffusion-based methods, yet remains tractable for networks up to *n* = 6000.

## V. Real-DATA VALIDATION

### A. Benchmark configuration

We benchmarked SPSID on the four gene–expression compendia released with the DREAM5 Network-Inference Challenge [22]. Table IV summarises the data sets: one simulated *in silico* network (Network 1) and three *in vivo* micro-array studies for *Staphylococcus aureus, Escherichia coli*, and *Saccharomyces cerevisiae*. Sample sizes range from 160 to 805, while the proportion of validated TF → Target edges (“Pos. ratio”) never exceeds 1.3 %, illustrating the severe class imbalance typical of GRN inference.

**TABLE IV:**
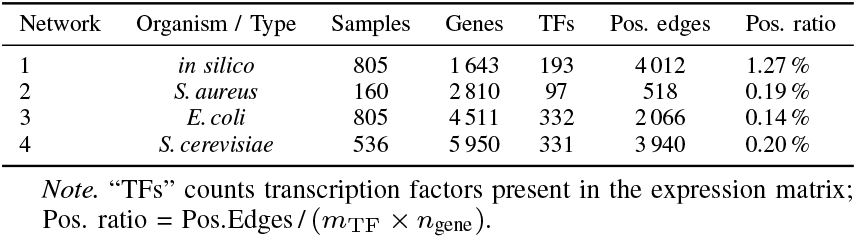
DREAM5 benchmark networks and their gold-standard statistics used in this study.

For each network we derived six raw score matrices with (i) Pearson correlation [23], (ii) Spearman correlation [23], (iii) GRNBoost2 [27], (iv) GENIE3 [26], (v) TIGRESS [25], and (vi) ARACNE [23], resulting in 4 × 6 = 24 upstream inference networks. Then each inference network was post-processed by our proposed SPSID and five baselines: RENDOR [10], ND [8], NE [9], Silencer [20], and ICM [49].

### B. Overall performance

We evaluated SPSID against five baseline denoising methods across the 24 DREAM5 benchmark tasks (4 networks × 6 upstream methods). The results are summarized in Fig. 4, which displays the AUROC (left) and AUPR (right) for every combination of network, inference engine, and denoising filter. Across both metrics, SPSID consistently yields the highest scores among all denoisers, indicated by the brightest cells in the heat-maps. For instance, when applying ARACNE to Network 1, SPSID achieves an AUPR of 0.197, which is substantially higher than the next-best denoiser, ND (0.191), and over six times better than RENDOR (0.031). This superior performance is consistent across the vast majority of tasks. A notable exception was on Network 1 with GENIE3 as the upstream method, where RENDOR’s scores (AUROC=0.973, AUPR=0.324) were marginally higher than those of SPSID (AUROC=0.968, AUPR=0.294). Furthermore, to address the known class-imbalance challenge, we also computed the Matthews Correlation Coefficient (MCC)[50], a more balanced metric. The full results for this metric are presented in **Supplementary Fig. S1**. This new heatmap confirms the findings from Fig. 4. SPSID is again the top-performing denoiser in the vast majority of cases. For instance, on the TIGRESS backbone (Supplementary Fig. S1E), SPSID achieves the highest MCC score on Network 1 (0.197), clearly outperforming the next-best denoisers, ND (0.118) and NE (0.087). Fig. 5 aggregates these results to show the score distributions for the raw network scores (Base) and the six denoising methods across all 24 tasks. The box plot for SPSID is distinctly superior for both AUROC (A) and AUPR (B), achieving the highest mean and median scores. The AUPR results of SPSID are higher than any baseline, and its lower quartile exceeds the median performance of all other methods. Conversely, methods such as Silencer and ICM often underperform relative to the raw network scores, suggesting they are less effective for these complex real-world datasets.

**Fig. 4:**
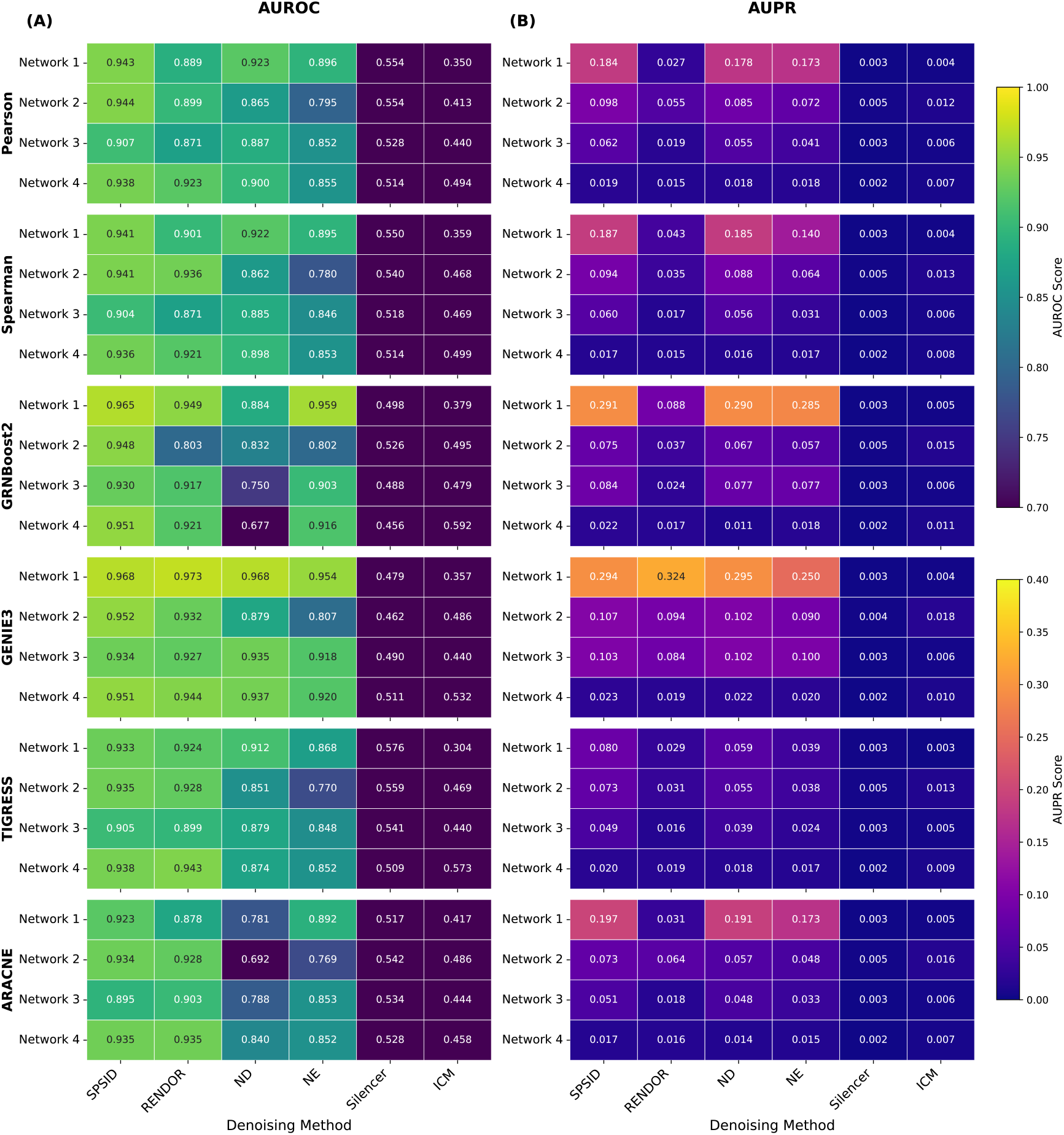
Facet heat-map of AUROC (left) and AUPR (right) across the four DREAM5 networks. Rows correspond to upstream inference back-bones (Pearson, Spearman, GENIE3, GRNBoost2, TIGRESS, and ARACNE); columns list the six denoising strategies examined. Colour intensity encodes metric value; exact scores are printed in each cell. Methods shown are SPSID (Single-Parameter Shrinkage Inverse-Diffusion), Reverse Network Diffusion On Random Walks (RENDOR), Network Deconvolution (ND), Network Enhancement (NE), Global Silencing of Indirect Correlations (Silencer), and Inverse Correlation Matrix (ICM).

**Fig. 5:**
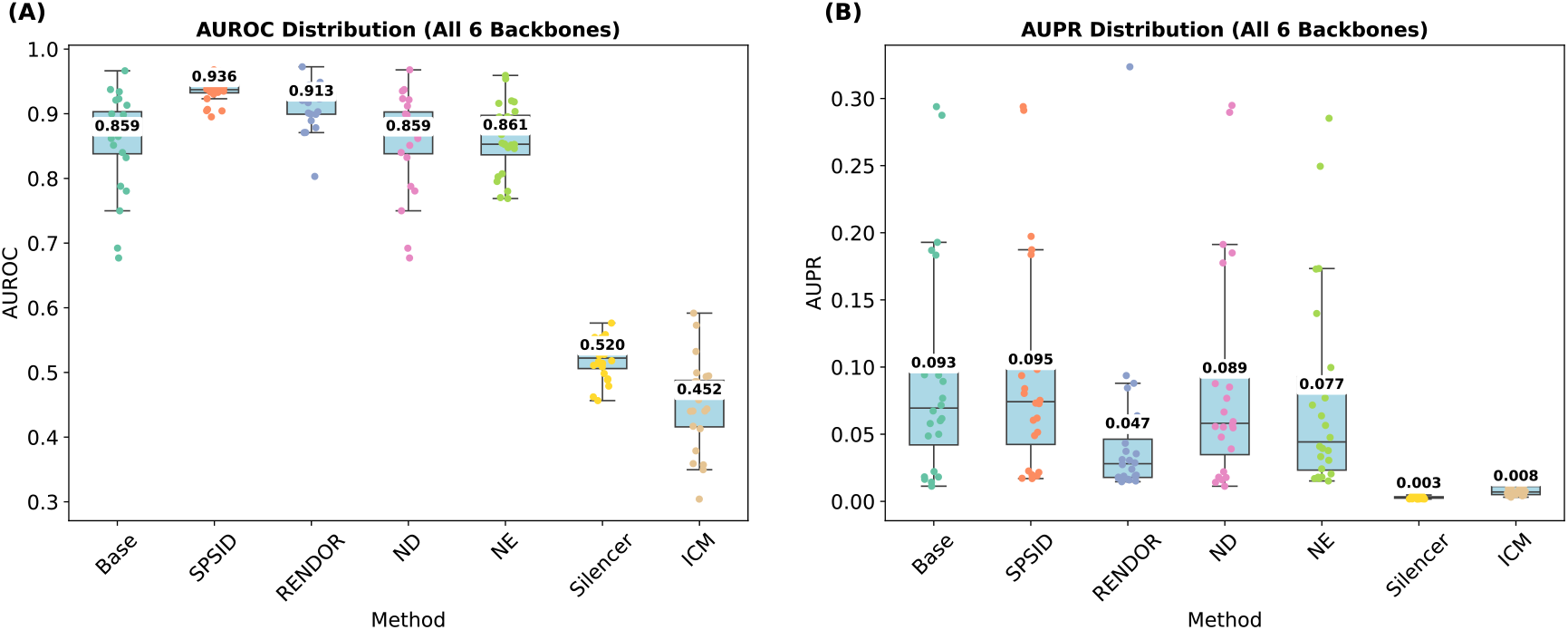
Box plots detailing the performance distribution over the 24 DREAM5 combinations (4 networks × 6 upstream inference methods) for the base network and each denoising method. (A) AUROC distribution. (B) AUPR distribution. For each method, the light blue box indicates the interquartile range (IQR) from the 25th to the 75th percentile, with the median marked by a horizontal line inside. The whiskers extend to show the data range. Coloured dots represent the individual scores for each task, and the arithmetic mean is printed as a numeric value with a light background. Methods shown are SPSID (Single-Parameter Shrinkage Inverse-Diffusion), Reverse Network Diffusion On Random Walks (RENDOR), Network Deconvolution (ND), Network Enhancement (NE), Global Silencing of Indirect Correlations (Silencer), and Inverse Correlation Matrix (ICM).

### C. Performance gains of denoising filters on DREAM5 inferred networks

To quantify how effectively each denoising filter improves upon the upstream inference networks, we calculated the percentage improvement in AUROC and AUPR relative to the raw network scores (Base) across the 24 DREAM5 tasks (4 networks × 6 upstream methods). Table V summarises the median and interquartile range (IQR) of these improvements. The results clearly show that SPSID is the only method to achieve a positive median improvement in both AUROC (+7.81 %) and AUPR (+3.29 %). RENDOR also delivers a positive median improvement in AUROC (+3.30 %) but its median AUPR change is strongly negative (− 50.97 %), reflecting the method’s occasional tendency to distort precision when positives are extremely scarce. ND hovers near zero for AUROC (0.00 %) and has a slightly negative AUPR median (− 1.62 %), indicating near–neutral impact. Network Enhancement, Silencer and ICM exhibit negative medians throughout; in particular, Silencer and ICM decrease AUPR by more than 87 percentage points, confirming that wholesale spectral suppression without shrinkage can be detrimental under class imbalance.

**TABLE V:** Overall percentage improvement over the raw (Base) network across all 24 DREAM5 tasks (4 networks × 6 upstream methods), reported as median and Interquartile Range.

| Method | AUROC (%) |  | AUPR (%) |  |
| --- | --- | --- | --- | --- |
|  | Median | IQR | Median | IQR |
| SPSID | 7.81 | 9.64 | 3.29 | 6.03 |
| RENDOR | 3.30 | 9.28 | -50.97 | 59.25 |
| ND | 0.00 | 0.05 | -1.62 | 3.45 |
| NE | -2.91 | 8.09 | -13.34 | 26.58 |
| Silencer | -39.24 | 7.15 | -94.95 | 4.81 |
| ICM | -45.62 | 8.83 | -87.60 | 25.60 |

*Note.* Values are *median* and *Interquartile Range (IQR)*; IQR is the difference between the 75th and 25th percentiles (Q3 - Q1) over the six upstream inference methods (Pearson, Spearman, GENIE3, GRNBoost2, TIGRESS, and ARACNE). Methods shown are SPSID (Single-Parameter Shrinkage Inverse-Diffusion), Reverse Network Diffusion On Random Walks (RENDOR), Network Deconvolution (ND), Network Enhancement (NE), Global Silencing of Indirect Correlations (Silencer), and Inverse Correlation Matrix (ICM).

Decomposing the performance gains by each of the four DREAM5 networks (Table VI) reveals that SPSID’s advantage is universal. It is the only method to achieve a positive median increase in both metrics across all four networks, with particularly strong gains on Network 2 (*S. aureus*) (9.53% in AUROC, 4.73% in AUPR). On the easiest *in silico* network (Network 1) it produces a small but consistent AUROC lift (2.16%) and keeps AUPR essentially unchanged (0.21%), demonstrating that it does not over-correct when indirect noise is already minimal. On the three *in vivo* data sets the gains are markedly larger. For *S. aureus* (Network 2) SPSID adds over nine percentage points (9.53%) in AUROC and nearly five in AUPR (4.73%), whereas RENDOR improves AUROC by over seven points (7.32%) but simultaneously reduces AUPR by more than forty percentage points (− 42.89%). Comparable patterns appear in *E. coli* (Network 3) and *S. cerevisiae* (Network 4), where SPSID remains the only method that increases both metrics simultaneously; all other filters either improve AUROC at the expense of AUPR or degrade both.

**TABLE VI:** Percentage improvement for each DREAM5 network, reported as median and Interquartile Range (IQR).

| Method | Network 1 (*in silico*) |  |  |  | Network 2 (*S. aureus*) |  |  |  |
| --- | --- | --- | --- | --- | --- | --- | --- | --- |
|  | AUROC<sub>med</sub> | AUROC<sub>IQR</sub> | AUPR<sub>med</sub> | AUPR<sub>IQR</sub> | AUROC<sub>med</sub> | AUROC<sub>IQR</sub> | AUPR<sub>med</sub> | AUPR<sub>IQR</sub> |
| ICM | -61.57 | 4.72 | -97.65 | 0.58 | -44.84 | 3.91 | -82.39 | 5.45 |
| ND | 0.04 | 0.05 | -0.93 | 2.68 | 0.01 | 0.05 | -1.87 | 6.26 |
| NE | -2.06 | 9.06 | -12.71 | 15.98 | -8.10 | 4.46 | -20.80 | 10.97 |
| RENDOR | 0.90 | 7.30 | -73.20 | 12.68 | 7.32 | 4.41 | -42.89 | 37.61 |
| SPSID | 2.16 | 5.35 | 0.21 | 0.96 | 9.53 | 3.69 | 4.73 | 5.46 |
| Silencer | -40.12 | 5.09 | -98.51 | 0.55 | -36.33 | 2.39 | -94.13 | 1.66 |

| Method | Network 3 (*E. coli*) |  |  |  | Network 4 (*S. cerevisiae*) |  |  |  |
| --- | --- | --- | --- | --- | --- | --- | --- | --- |
|  | AUROC<sub>med</sub> | AUROC<sub>IQR</sub> | AUPR<sub>med</sub> | AUPR<sub>IQR</sub> | AUROC<sub>med</sub> | AUROC<sub>IQR</sub> | AUPR<sub>med</sub> | AUPR<sub>IQR</sub> |
| ICM | -48.46 | 5.73 | -90.15 | 2.28 | -43.81 | 8.33 | -50.63 | 7.40 |
| ND | 0.01 | 0.04 | -4.43 | 8.27 | -0.01 | 0.01 | -1.15 | 1.24 |
| NE | -2.61 | 9.59 | -32.60 | 34.66 | -2.17 | 4.98 | 1.36 | 10.67 |
| RENDOR | 0.78 | 12.88 | -67.95 | 5.24 | 5.24 | 7.91 | -3.77 | 21.25 |
| SPSID | 2.60 | 8.70 | 0.58 | 4.44 | 5.78 | 6.07 | 6.73 | 12.00 |
| Silencer | -39.48 | 5.41 | -95.01 | 1.91 | -42.29 | 4.56 | -89.64 | 2.59 |

*Note.* Values are *median* and *Interquartile Range (IQR)*; IQR is the difference between the 75th and 25th percentiles (Q3 - Q1) over the six upstream inference methods. Methods shown are SPSID (Single-Parameter Shrinkage Inverse-Diffusion), Reverse Network Diffusion On Random Walks (RENDOR), Network Deconvolution (ND), Network Enhancement (NE), Global Silencing of Indirect Correlations (Silencer), and Inverse Correlation Matrix (ICM).

These numerical trends are mirrored by the diverging heatmaps in Fig. 6. Panel A shows that SPSID produces warm (red) tiles across almost the entire grid for AUROC, with particularly intense shades when used after GRNBoost2, and also showing consistent gains for the new ARACNE and TIGRESS backbones. Panel B confirms a similar, though more heterogeneous, pattern for AUPR: SPSID maintains positive values in twenty out of twenty-four combinations, while every baseline displays extensive blue regions indicating performance loss. To evaluate the ability of each method to jointly improve both metrics, we plotted the absolute improvements (ΔAUROC vs. ΔAUPR) in a scatter plot (Fig. 6C). The result is unambiguous: nearly all data points for SPSID are concentrated in the top-right quadrant, representing a simultaneous improvement in both AUROC and AUPR. In stark contrast, points for all other baseline methods are scattered across the other three quadrants, particularly the bottom-right (improving AUROC at the expense of AUPR, as seen with RENDOR) and the bottom-left (degrading both metrics, as with Silencer and ICM). Collectively, the median-IQR statistics and the heat-map visualisation demonstrate that the shrinkage inverse-diffusion strategy implemented in SPSID consistently enhances discriminative power and class-imbalance precision across a wide spectrum of networks and upstream inference engines, whereas competing filters either provide limited benefit or, in many cases, undermine baseline performance.

**Fig. 6:**
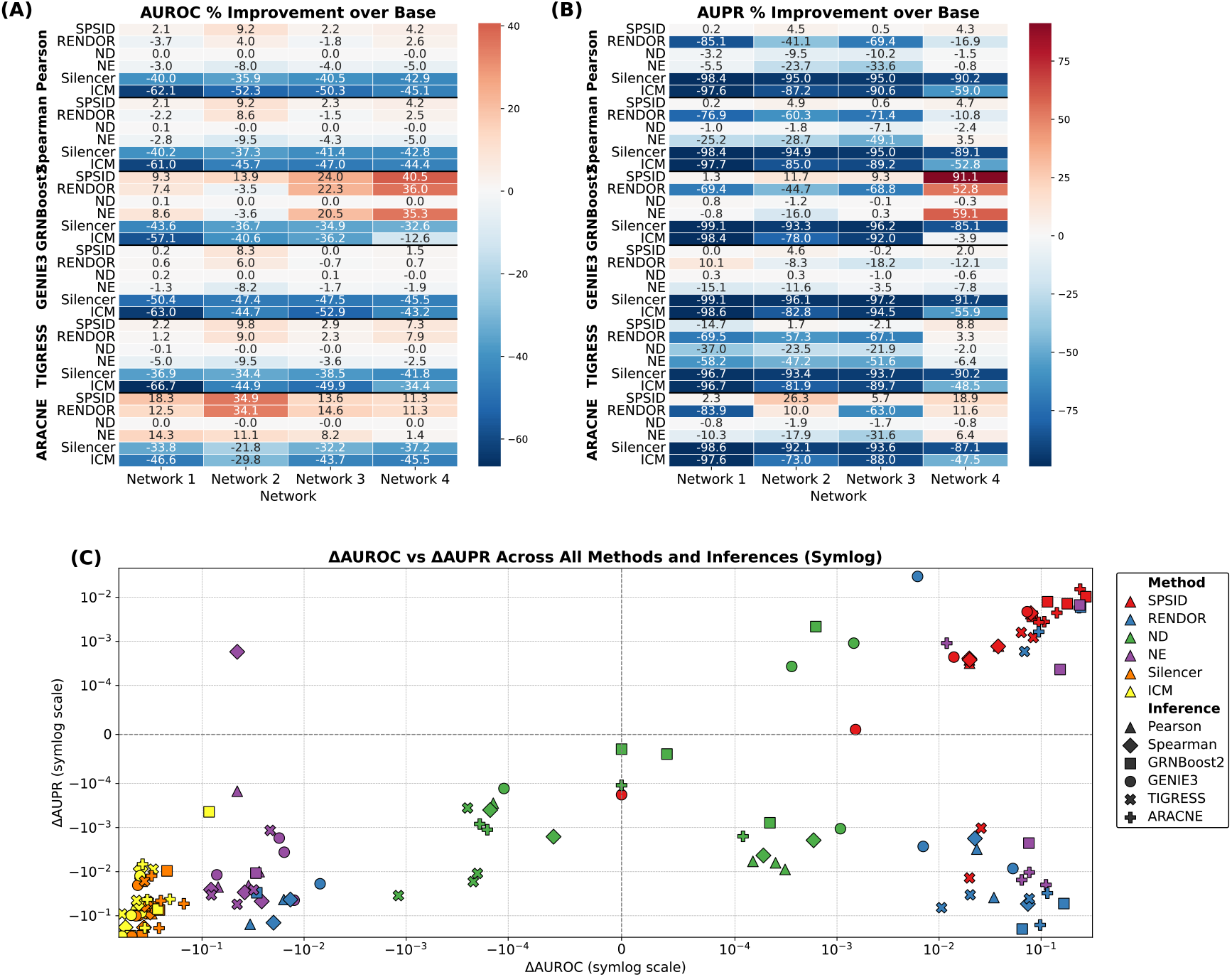
Performance improvement of denoising methods over the base network across the 24 DREAM5 tasks. (A) and (B) diverging heatmaps show the per-task percentage improvement for AUROC and AUPR, respectively. Red indicates an improvement over the baseline, while blue indicates a decrease. (C) A scatter plot showing the joint distribution of absolute improvements (ΔAUROC vs. ΔAUPR) on a symmetric-logarithmic scale. The color of each point corresponds to the denoising method, and the marker shape represents the upstream inference method. The denoising methods shown are SPSID (Single-Parameter Shrinkage Inverse-Diffusion), Reverse Network Diffusion On Random Walks (RENDOR), Network Deconvolution (ND), Network Enhancement (NE), Global Silencing of Indirect Correlations (Silencer), and Inverse Correlation Matrix (ICM).

### D. Rank-score comparison across methods

To assess performance consistency beyond absolute scores, we adopted a rank-based evaluation. For each of the 24 DREAM5 combinations (4 networks × 6 upstream inference methods), we ranked the *M* competing methods based on their AUROC and AUPR scores.

The rank *r*_*i*_ ∈ {1,…,*M}* for a denoising method *i* on a given task is converted to a Rank-score (*s*_*i*_) as follows:

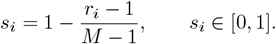

This score normalises the rank to the [0, 1] interval, where the top-performing method receives a score of 1 and the worst receives a score of 0. This approach allows for robust aggregation of relative performance across different tasks.

Fig. 7 provides a comprehensive analysis of these ranks. Panels A and B show that SPSID attains nearly perfect mean rank-scores in both metrics and displays the narrowest inter-quartile bands, confirming that it never falls below second place for any network–inference combination. RENDOR and ND form a middle tier whose mean scores are 0.15–0.25 points lower and whose variability is appreciably larger, indicating sensitivity to the choice of upstream engine. Network Enhancement trails this pair, whereas Silencer and ICM cluster close to zero, often occupying the last rank when precision-recall performance is considered. Panels C and D provide the Friedman average ranks: SPSID achieves a near-perfect 1.17 for AUROC and 1.08 for AUPR, versus 2.25 (RENDOR) and 2.38 (ND) for the nearest competitors in each metric, respectively. The overall Friedman statistics are now even more significant (*χ*^2^ = 106.19 for AUROC, *χ*^2^ = 108.19 for AUPR) and correspond to *p* = 2.61e − 21 and *p* = 9.88e − 22 respectively, decisively rejecting the null hypothesis that all filters perform equivalently. These rank-based assessments, which are independent of absolute metric magnitude, reinforce earlier percentage-gain results: SPSID not only delivers the good average accuracy but also the consistent behaviour across disparate data sets and upstream inference pipelines.

**Fig. 7:**
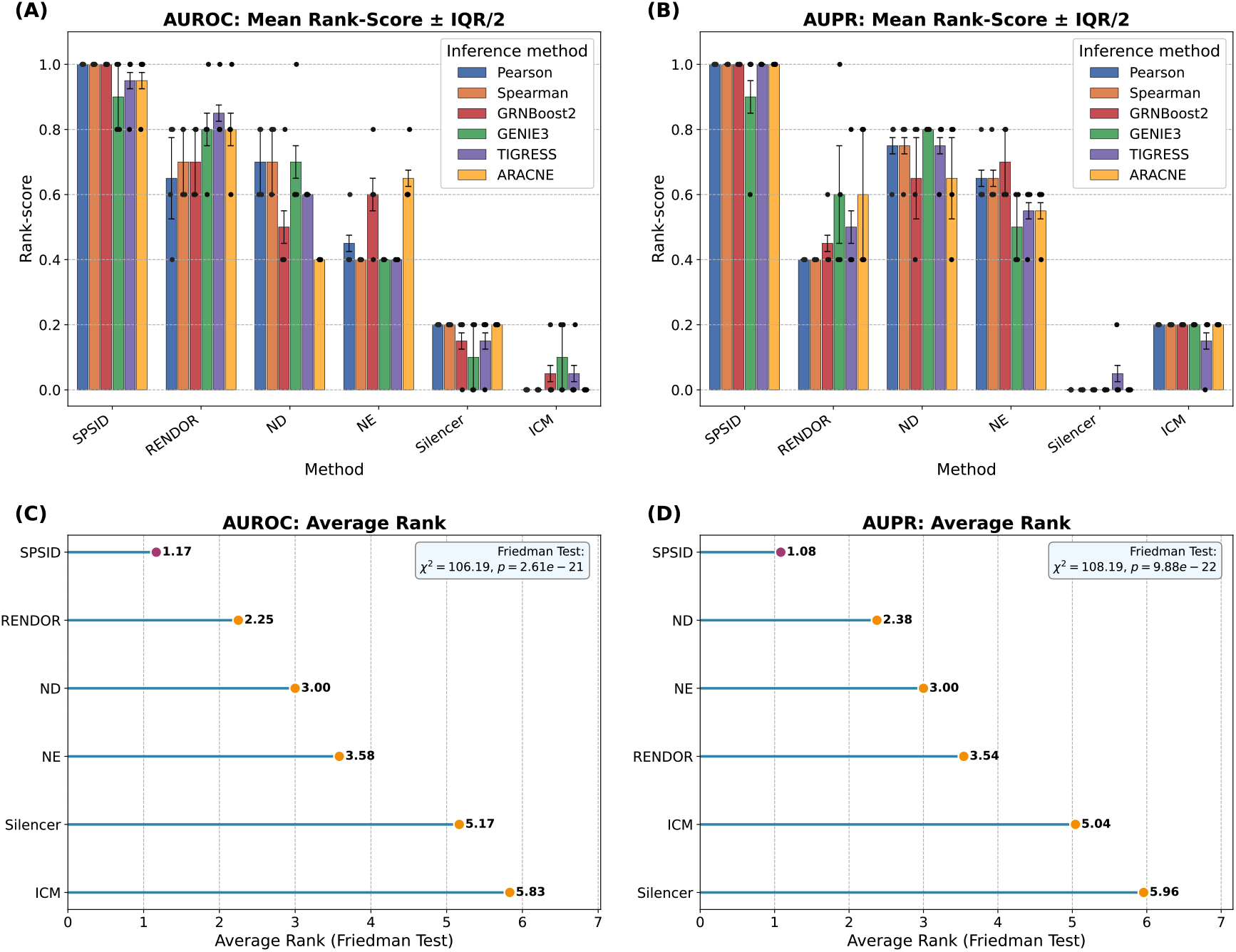
A comprehensive analysis of method ranks over the 24 DREAM5 combinations. (A) and (B) Grouped bar charts display the mean rank-score for AUROC and AUPR, respectively. Error bars indicate half the inter-quartile range (IQR/2), providing a measure of variance across the six inference engines. (C) and (D) Lollipop plots show the average ordinal rank determined by the Friedman test for AUROC and AUPR. In all panels, superior performance is indicated by a higher rank-score (for the bar charts) or a lower average rank (for the lollipop plots). Methods shown are SPSID (Single-Parameter Shrinkage Inverse-Diffusion), Reverse Network Diffusion On Random Walks (RENDOR), Network Deconvolution (ND), Network Enhancement (NE), Global Silencing of Indirect Correlations (Silencer), and Inverse Correlation Matrix (ICM).

### E. Validation on scRNA-seq benchmarks

To further validate the robustness and generalizability of SPSID, we evaluated it on seven scRNA-seq datasets curated by the BEELINE benchmark [51]: mouse embryonic stem cells (mESC; GEO: GSE98664) [52], mouse hematopoietic stem/progenitor cells with three lineage subsets (mHSC-E/GM/L; GEO: GSE81682) [53], mouse dendritic cells (mDC; GEO: GSE48968) [54], human embryonic stem cells (hESC; GEO: GSE75748) [55], and human mature hepatocytes (hHEP; GEO: GSE81252) [56]. Following BEELINE’s preprocessing protocol [51], we applied all six upstream inference methods (Pearson, Spearman, GENIE3, GRNBoost2, TIGRESS, and ARACNE), followed by SPSID and all five competing denoisers. We evaluated all results against the cell-type-specific ground truth network. The full results are presented in **Supplementary Fig. S2**. The results are conclusive: SPSID was the top-performing denoising method in the vast majority of the 42 combinations (7 datasets × 6 upstream methods), achieving the highest AUROC in 37 out of 42 (88%) tasks and the highest AUPR in 34 out of 42 (81%) tasks. For instance, on the hESC-GRNBoost2 task, SPSID achieved top-performing scores for both AUROC (0.895) and AUPR (0.029). This comprehensive benchmark confirms that SPSID’s denoising capability is highly effective and robust, generalizing successfully to sparse single-cell data.

### F. Sensitivity to GRN incompleteness

To assess the robustness of SPSID to incompleteness in current GRN annotations, we exploited the historical snapshots provided by Abasy Atlas v2.2 for *E. coli* K-12 MG1655 (NCBI 511145) [57]. Abasy consolidates and meta-curates bacterial regulatory networks from multiple sources, and its updated release provides snapshots at different curation years, together with estimates of network completeness and systems-level features such as feed-forward (FF) motifs.

Using the official mapping file for DREAM5 Network 3, we mapped all 1081 genes (141 TFs and 999 targets) to their corresponding Abasy gene names. For each of the nine available snapshots (2003, 2005, 2006, 2011, 2013, 2014, 2015, 2017, 2018) we extracted all TF–target interactions whose endpoints belonged to this gene set, obtaining a series of gold standards with increasing completeness (from approximately 1.2 × 10^3^ to 5.5 × 10^3^ edges). The upstream inferred networks were kept fixed and identical to our DREAM5 experiments (Pearson, Spearman, GRNBoost2, GENIE3, TIGRESS, ARACNE). For each (backbone, snapshot) pair, we applied SPSID and all competing denoisers and evaluated AUROC and AUPR.

Supplementary Fig. S3 shows the resulting AUPR trajectories as a function of snapshot year. As the Abasy gold standard becomes more complete, AUPR decreases for all methods and backbones, reflecting that later snapshots incorporate more condition-specific and weaker interactions that are harder to recover. Crucially, the curves for different denoisers are almost parallel: SPSID remains the best or tied-best method at all completeness levels, and its advantage over NE and ND is essentially constant across years. Supplementary Fig. S4 reports the corresponding AUROC results, which are relatively stable or slightly increasing with completeness, again with SPSID consistently at or near the top.

These results indicate that gold-standard incompleteness primarily affects the absolute level of the evaluation metrics (earlier snapshots leading to more optimistic AUPR), but it does not qualitatively change the ranking among denoising methods. Moreover, although the number of FF motifs increases with network completeness in Abasy [57], we do not observe any deterioration in SPSID’s relative performance at later snapshots. Thus, SPSID does not appear to over-penalize true FF-mediated transitive interactions; instead, it provides robust gains across a wide range of GRN completeness levels.

## VI. Discussion

Inferring gene regulatory networks from high-dimensional expression data is a central challenge in computational biology, complicated by both experimental noise and, more critically, structural noise arising from transitive correlations [20]. In this work, we introduced SPSID, a shrinkage inverse-diffusion operator designed to specifically address this latter challenge. Our comprehensive evaluations on both simulated data and the gold-standard DREAM5 benchmark demonstrate that SPSID achieves good accuracy and exhibits robustness.

The primary innovation of SPSID lies in its elegant integration of a Tikhonov shrinkage regulariser (*λI*) into a network deconvolution framework. This single modification provides a dual benefit that underpins its superior performance. First, it guarantees the numerical stability of the matrix inversion, a known vulnerability in classical deconvolution methods that may require ad-hoc data scaling to ensure the convergence of the Neumann series [8]. Second, and more importantly, the shrinkage term provides a principled mechanism for differentially attenuating network signals based on their path length. As shown in our analysis (Section III-B), the operator strongly suppresses eigenvalues associated with multi-step, indirect paths while preserving the relative strength of signals corresponding to direct, one-step interactions. This allows SPSID to effectively function as a low-pass spectral filter that isolates direct regulatory links. This design contrasts with other diffusion-based methods like Network Enhancement [9] or RENDOR [10], which, despite their effectiveness, may require tuning of data-dependent hyperparameters to balance the diffusion process.

The empirical results presented in this study validate our theoretical framework. In simulations, SPSID consistently outperformed five competing methods across diverse conditions of network sparsity, noise levels, and transitive effect strengths (Section IV). This robustness is critical, as these underlying network properties are typically unknown in real-world applications. On the DREAM5 benchmark [22], four networks processed by six heterogeneous inference engines (Pearson and Spearman, GRNBoost2, GENIE3, TIGRESS, and ARACNE), SPSID improves both AUROC and AUPR in 20 out of 24 combinations. ND and RENDOR raise AUROC in some cases but often decrease AUPR, particularly for extremely imbalanced *in vivo* data sets, confirming that shrinkage is essential when precision–recall performance is the primary concern. Global–silencing approaches such as Silencer [20] and ICM [49] frequently degrade both metrics, supporting long-standing concerns that indiscriminate spectral suppression destroys true signal. Furthermore, the DREAM5 results are characterized by a severe class imbalance (e.g., 0.14% positives in E. coli), which results in low absolute AUPR values for all methods. It is important to frame performance in this context, and to look beyond aggregate medians. As shown in the per-task heatmap (Fig. 6B), ‘marginal’ median gains (like 0.58% on Net 3) coexist with massive, non-marginal gains on specific backbones (e.g., +24.0% on Net 3 with GRNBoost2). This also explains the performance difference between networks: E. coli (Net 3) is the most challenging benchmark, with the highest sparsity (0.14% positives) and faintest signal. The modest gains on this network reflect the fundamental difficulty of this specific biological dataset. More importantly, SPSID is the only method to provide consistent positive gains, whereas competing filters like RENDOR are highly unstable. To provide a more balanced assessment in this imbalanced context, MCC results further support our findings, showing that these modest AUPR gains translate to clear and robust performance increases in MCC and confirming the practical utility of our method.

A valid observation from our benchmark is that the performance gains from SPSID are more modest when applied to already-sophisticated backbones like GENIE3, compared to the massive gains on simpler methods like Pearson correlation. This is expected, as GENIE3 itself is a powerful feature selection algorithm. However, the true utility of SPSID is demonstrated by its robustness: unlike other filters that can be unstable and catastrophically degrade strong networks, SPSID provides a stable, positive (albeit incremental) gain. This “do no harm” property, combined with its parameter-free nature, makes it a uniquely safe and reliable general-purpose tool, in stark contrast to the significant improvement in stability it offers over tuning-sensitive methods.

It is important to clarify the biological interpretation of SPSID’s output. Upstream inference engines often capture all strong statistical associations, confounding direct causal links (A → B) with indirect transitive correlations (A → B → C, creating a spurious A-C link). SPSID is designed to deconvolve these pathways and filter out the latter. Consequently, an interaction that retains a high score after SPSID processing should be interpreted as a high-confidence candidate for a direct regulatory interaction, such as the physical binding of a transcription factor to its target’s regulatory element. For experimentalists, this provides a more reliable and prioritized list of interactions for downstream validation, reducing the risk of pursuing false positives derived from multi-step network effects. While we have focused on GRNs, the problem of disentangling direct from indirect effects is pervasive across systems biology. The principles underlying SPSID are general and could readily be applied to other types of interaction networks, such as protein-protein interaction (PPI) networks derived from high-throughput screens [58], metabolic networks reconstructed from flux balance analysis [59], or even ecological interaction webs. Any system where interactions can be represented as a weighted adjacency matrix plagued by transitive effects is a potential candidate for SPSID-based denoising.

A key biological challenge is the Feed-Forward Loop (FFL) motif, where a direct link (A → C) co-exists with an indirect path (A → B → C). Our deconvolution framework is ideally suited for this. The observed A-C score is a mixture of the 1-step *P*_dir_(*A, C*) signal and the 2-step 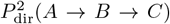 noise. SPSID is designed to subtract the 2-step noise component, thereby *isolating* the true *P*_dir_(*A, C*) direct link, not eliminating it. In contrast, for a purely spurious link (A → D) that only exists via 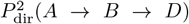, the subtraction correctly results in a near-zero score. This act of ‘cleaning’ the network, rather than just ‘silencing’ it, makes the SPSID output a more reliable substrate for downstream motif analysis.

Looking forward, several avenues for future research are apparent. A key direction is the adaptation of SPSID to the unique challenges of single-cell RNA-sequencing (scRNA-seq) data. The extreme sparsity (dropout events) and stochastic nature of single-cell data, where noise can play functional roles, require specialized modeling approaches [60], [61]. Extending the SPSID framework to handle dynamic, time-course data or to integrate multi-omics information could further enhance its power and applicability. For instance, incorporating information from ATAC-seq to inform the prior probability of regulatory links could create a more powerful, integrated inference pipeline. Another valuable direction would be to integrate SPSID into an ensemble framework. For instance, applying SPSID as a post-processing step to a set of networks generated by diverse upstream engines (e.g., GENIE3, TIGRESS, ARACNE) and then creating a consensus network could further enhance robustness and accuracy.

A key strength of SPSID is its robustness to the shrinkage parameter *λ*, which allows for the use of a fixed default and removes the need for data-dependent tuning. Our analysis confirms that performance is virtually identical for any large *λ*. Nonetheless, a valid question is whether an adaptive parameter could yield better results in extreme network conditions. While this would re-introduce the risk of overfitting that our method was designed to avoid, exploring principled, adaptive shrinkage methods (e.g., based on network topology or noise estimates) remains a valuable direction for future research, especially when applying the method to networks with fundamentally different structures.

Another practical consideration is scalability. Our current implementation uses direct matrix inversion, which scales at *O*(*n*^3^). As shown in our runtime analysis, this is computationally tractable and efficient for networks up to *n* = 6000, where SPSID still runs in a couple of minutes on a standard desktop. However, for very large-scale networks (e.g., *n>* 20,000), this approach becomes a bottleneck. A clear path forward is to avoid forming the inverse and instead solve the linear system

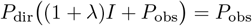

with iterative Krylov methods such as the Generalized Minimal Residual (GMRES) algorithm [62], [63] and the Biconjugate Gradient Stabilized (BiCGSTAB) method [64]. These solvers reduce the dominant cost to matrix–vector products (and preconditioning), thereby removing the explicit *O*(*n*^3^) step and paving the way for applying SPSID to much larger biological networks.

In conclusion, SPSID provides an effective and computationally efficient solution to the persistent problem of structural noise in biological network inference. By leveraging a principled shrinkage-based operator, it successfully distinguishes direct interactions from transitive correlations without the need for parameter tuning. The method’s demonstrated potential to enhance the performance of existing inference approaches suggests its utility as a valuable tool for researchers aiming to better elucidate the regulatory architecture of complex biological systems.

## VII. Conclusion

High–dimensional gene–regulatory networks inferred from omics data are severely contaminated by structural noise—indirect statistical dependencies that masquerade as direct regulatory links. In this paper, we addressed the persistent challenge of structural noise in gene regulatory network inference, a problem that often leads to a high number of spurious interactions due to transitive correlations. We introduced SPSID, a novel network denoising framework built upon a shrinkage inverse-diffusion operator. The core of our method is a mathematically principled approach that stabilizes the deconvolution process and systematically filters out indirect, multi-step network paths, thereby isolating direct regulatory signals.

Our extensive evaluations demonstrated the effectiveness and robustness of SPSID. Through comprehensive simulations, we showed that SPSID consistently outperforms existing state-of-the-art methods across a wide spectrum of network topologies, noise levels, and transitive effect strengths. In addition, when applied to the gold-standard DREAM5 benchmark, SPSID acted as a post-processing tool, significantly enhancing the accuracy of multiple upstream inference algorithms.

By providing a reliable method to distinguish direct from indirect interactions, our work offers a practical step towards mapping the regulatory architecture of complex biological systems with greater fidelity. The principles underlying SPSID are general, suggesting its potential utility for denoising other types of biological networks where transitive effects are a confounding factor.

## Data and Code Availability

The DREAM5 benchmark datasets are publicly available from the Synapse platform (https://www.synapse.org/).

## Conflict of interest

The authors declare that they have no competing interests.

## Supplementary Materials

**Table S1:** Runtime comparison on synthetic networks of increasing size *n*.

| Method \ $n$ | 500 | 1000 | 2000 | 3000 | 4000 | 5000 | 6000 |
| --- | --- | --- | --- | --- | --- | --- | --- |
| SPSID | $0.24 \pm 0.02$ | $1.01 \pm 0.02$ | $5.33 \pm 0.01$ | $17.49 \pm 0.17$ | $48.47 \pm 0.06$ | $89.67 \pm 0.14$ | $148.72 \pm 0.25$ |
| RENDOR | $0.14 \pm 0.03$ | $0.53 \pm 0.01$ | $5.04 \pm 0.16$ | $18.48 \pm 0.55$ | $53.94 \pm 0.21$ | $102.03 \pm 0.41$ | $174.87 \pm 1.24$ |
| ND | $0.42 \pm 0.00$ | $3.38 \pm 0.04$ | $53.70 \pm 0.32$ | $246.55 \pm 0.73$ | $651.77 \pm 1.24$ | $1298.00 \pm 2.70$ | $2373.05 \pm 1.05$ |
| NE | $0.11 \pm 0.00$ | $0.48 \pm 0.03$ | $2.53 \pm 0.02$ | $10.38 \pm 0.14$ | $23.06 \pm 0.62$ | $42.23 \pm 0.29$ | $71.24 \pm 0.16$ |
| Silencer | $0.03 \pm 0.01$ | $0.17 \pm 0.03$ | $0.82 \pm 0.02$ | $3.07 \pm 0.12$ | $6.09 \pm 0.14$ | $10.59 \pm 0.13$ | $16.92 \pm 0.23$ |
| ICM | $0.03 \pm 0.00$ | $0.17 \pm 0.07$ | $0.58 \pm 0.03$ | $2.09 \pm 0.17$ | $3.86 \pm 0.17$ | $6.26 \pm 0.06$ | $10.16 \pm 0.06$ |

*Notes:* Entries report mean ± standard deviation of wall-clock time (in seconds) across repeated runs on the same hardware.

**Table S2:** Mean AUROC / AUPR under varying sparsity (*ρ*), noise (*σ*) and transitivity (*β*).

| (a) Varying network density ($\sigma = 0.5$, $\beta = 0.5$) | | | | | | | | | | |
| --- | --- | --- | --- | --- | --- | --- | --- | --- | --- | --- |
| Method | $\rho = 0.01$ | | $\rho = 0.03$ | | $\rho = 0.05$ | | $\rho = 0.07$ | | $\rho = 0.09$ | |
|  | AUROC | AUPR | AUROC | AUPR | AUROC | AUPR | AUROC | AUPR | AUROC | AUPR |
| SPSID | **0.909** | **0.177** | **0.896** | **0.277** | **0.878** | **0.305** | **0.856** | **0.313** | **0.832** | **0.317** |
| RENDOR | 0.865 | 0.096 | 0.842 | 0.170 | 0.816 | 0.203 | 0.789 | 0.222 | 0.764 | 0.238 |
| ND | 0.857 | 0.105 | 0.835 | 0.164 | 0.807 | 0.187 | 0.778 | 0.201 | 0.751 | 0.214 |
| NE | 0.830 | 0.089 | 0.810 | 0.168 | 0.777 | 0.199 | 0.744 | 0.216 | 0.717 | 0.230 |
| Silencer | 0.526 | 0.012 | 0.521 | 0.033 | 0.517 | 0.054 | 0.514 | 0.074 | 0.511 | 0.094 |
| ICM | 0.453 | 0.009 | 0.460 | 0.027 | 0.467 | 0.046 | 0.472 | 0.064 | 0.476 | 0.084 |

| (b) Varying noise level ($\rho = 0.05$, $\beta = 0.5$) | | | | | | | | | | |
| --- | --- | --- | --- | --- | --- | --- | --- | --- | --- | --- |
| Method | $\sigma = 0.1$ | | $\sigma = 0.3$ | | $\sigma = 0.5$ | | $\sigma = 0.7$ | | $\sigma = 0.9$ | |
|  | AUROC | AUPR | AUROC | AUPR | AUROC | AUPR | AUROC | AUPR | AUROC | AUPR |
| SPSID | **0.957** | **0.528** | **0.940** | **0.454** | **0.878** | **0.303** | **0.811** | **0.208** | **0.758** | **0.156** |
| RENDOR | 0.939 | 0.379 | 0.894 | 0.296 | 0.816 | 0.202 | 0.750 | 0.146 | 0.703 | 0.116 |
| ND | 0.925 | 0.332 | 0.887 | 0.270 | 0.807 | 0.187 | 0.740 | 0.135 | 0.693 | 0.108 |
| NE | 0.884 | 0.311 | 0.853 | 0.278 | 0.777 | 0.199 | 0.715 | 0.144 | 0.671 | 0.114 |
| Silencer | 0.537 | 0.059 | 0.523 | 0.055 | 0.516 | 0.054 | 0.514 | 0.053 | 0.511 | 0.052 |
| ICM | 0.439 | 0.042 | 0.455 | 0.044 | 0.466 | 0.045 | 0.473 | 0.046 | 0.478 | 0.047 |

| (c) Varying transitive strength ($\rho = 0.05$, $\sigma = 0.5$) | | | | | | | | | | |
| --- | --- | --- | --- | --- | --- | --- | --- | --- | --- | --- |
| Method | $\beta = 0.1$ | | $\beta = 0.3$ | | $\beta = 0.5$ | | $\beta = 0.7$ | | $\beta = 0.9$ | |
|  | AUROC | AUPR | AUROC | AUPR | AUROC | AUPR | AUROC | AUPR | AUROC | AUPR |
| SPSID | **0.909** | **0.446** | **0.897** | **0.381** | **0.879** | **0.304** | **0.858** | **0.237** | **0.837** | **0.188** |
| RENDOR | 0.855 | 0.269 | 0.839 | 0.240 | 0.816 | 0.201 | 0.792 | 0.166 | 0.765 | 0.136 |
| ND | 0.847 | 0.251 | 0.831 | 0.226 | 0.807 | 0.187 | 0.780 | 0.150 | 0.752 | 0.123 |
| NE | 0.804 | 0.245 | 0.796 | 0.227 | 0.777 | 0.199 | 0.754 | 0.168 | 0.725 | 0.141 |
| Silencer | 0.525 | 0.054 | 0.522 | 0.054 | 0.518 | 0.054 | 0.513 | 0.053 | 0.510 | 0.053 |
| ICM | 0.460 | 0.045 | 0.460 | 0.045 | 0.466 | 0.045 | 0.471 | 0.046 | 0.478 | 0.047 |

*Note.* Bold values indicate the best results. Methods shown are SP SID (Single-Parameter Shrinkage Inverse-Diffusion), Reverse Network Diffusion On Random Walks (RENDOR), Network Deconvolution (ND), Network Enhancement (NE), Global Silencing of Indirect Correlations (Silencer), and Inverse Correlation Matrix (ICM).

**Table S3:**
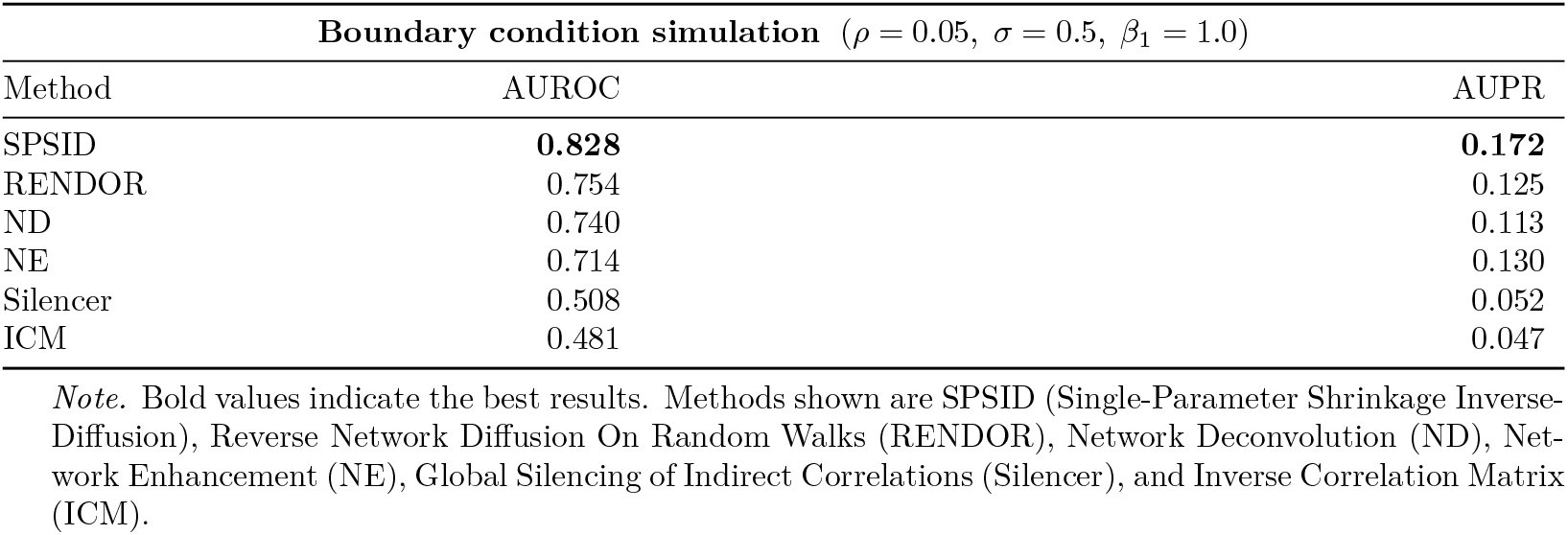
Mean AUROC / AUPR on G2 simulation model at boundary condition (*β*_1_ = 1.0).

| Boundary condition simulation ($\rho = 0.05, \sigma = 0.5, \beta_1 = 1.0$) | | |
| --- | --- | --- |
| Method | AUROC | AUPR |
| SPSID | **0.828** | **0.172** |
| RENDOR | 0.754 | 0.125 |
| ND | 0.740 | 0.113 |
| NE | 0.714 | 0.130 |
| Silencer | 0.508 | 0.052 |
| ICM | 0.481 | 0.047 |

*Note.* Bold values indicate the best results. Methods shown are SP SID (Single-Parameter Shrinkage Inverse-Diffusion), Reverse Network Diffusion On Random Walks (RENDOR), Network Deconvolution (ND), Network Enhancement (NE), Global Silencing of Indirect Correlations (Silencer), and Inverse Correlation Matrix (ICM).

**Table S4:** Mean AUROC / AUPR on higher-order simulation model.

| Varying cubic (G3) transitive strength ($\beta_2$) ($\rho = 0.05, \sigma = 0.5, \beta_1 = 0.5$) | | | | | | | | | | |
| --- | --- | --- | --- | --- | --- | --- | --- | --- | --- | --- |
| Method | $\beta_2 = 0.1$ | | $\beta_2 = 0.3$ | | $\beta_2 = 0.5$ | | $\beta_2 = 0.7$ | | $\beta_2 = 0.9$ | |
|  | AUROC | AUPR | AUROC | AUPR | AUROC | AUPR | AUROC | AUPR | AUROC | AUPR |
| SPSID | **0.869** | **0.271** | **0.829** | **0.202** | **0.788** | **0.157** | **0.757** | **0.131** | **0.733** | **0.115** |
| RENDOR | 0.814 | 0.201 | 0.785 | 0.178 | 0.750 | 0.147 | 0.719 | 0.123 | 0.694 | 0.108 |
| ND | 0.805 | 0.184 | 0.775 | 0.152 | 0.736 | 0.123 | 0.705 | 0.105 | 0.679 | 0.094 |
| NE | 0.764 | 0.189 | 0.727 | 0.158 | 0.693 | 0.133 | 0.667 | 0.115 | 0.649 | 0.103 |
| Silencer | 0.517 | 0.054 | 0.512 | 0.054 | 0.508 | 0.053 | 0.506 | 0.053 | 0.505 | 0.052 |
| ICM | 0.467 | 0.045 | 0.475 | 0.046 | 0.480 | 0.047 | 0.485 | 0.047 | 0.487 | 0.048 |

*Note.* Bold values indicate the best results. Methods shown are SP SID (Single-Parameter Shrinkage Inverse-Diffusion), Reverse Network Diffusion On Random Walks (RENDOR), Network Deconvolution (ND), Network Enhancement (NE), Global Silencing of Indirect Correlations (Silencer), and Inverse Correlation Matrix (ICM).

**Supplementary Fig. S1:**
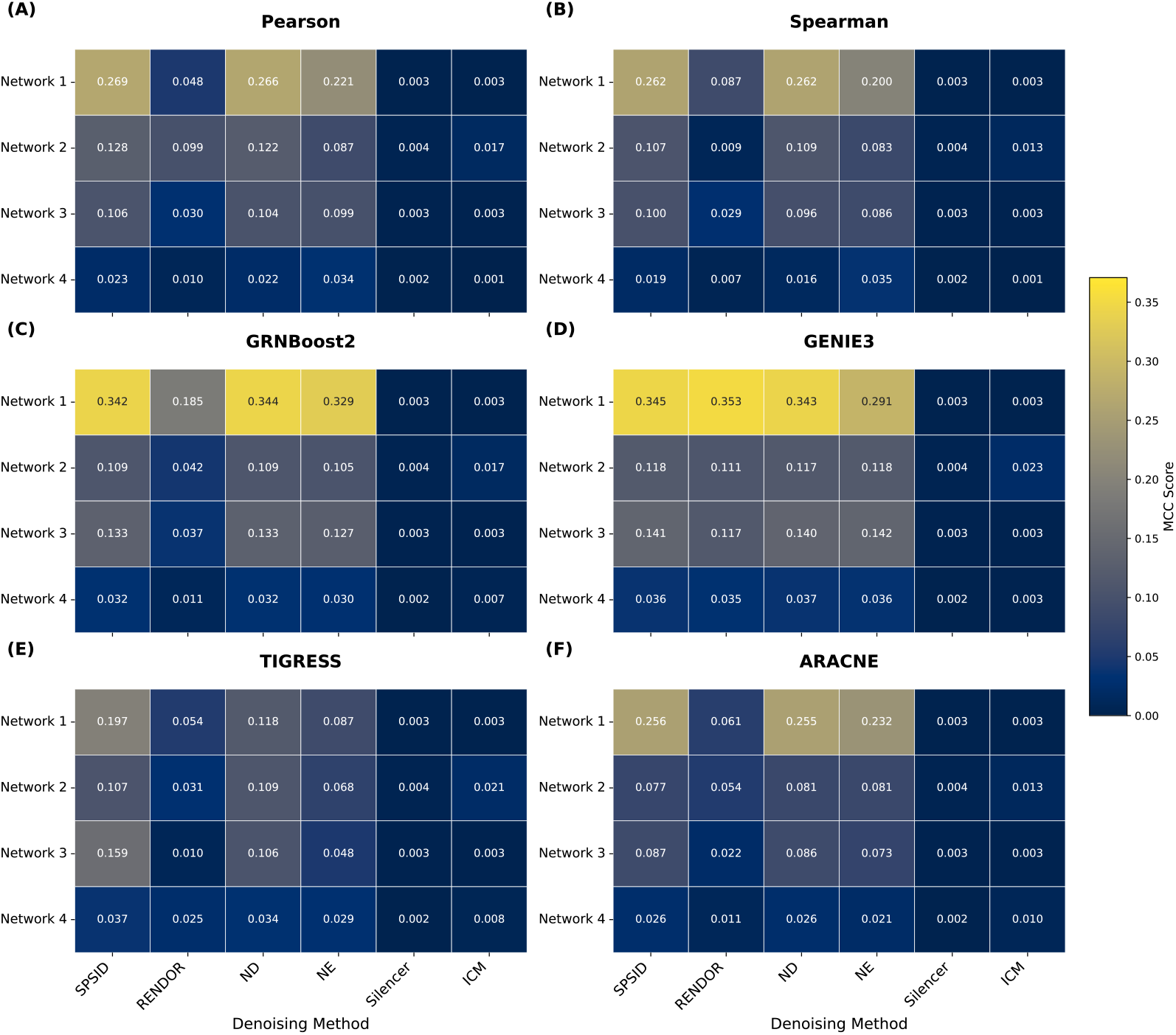
DREAM5 MCC benchmark performance. Facet heatmap displaying MCC scores. Each panel (A-F) represents one of the 6 upstream inference methods. Columns compare SPSID against 5 other denoising baselines across the 4 DREAM5 networks.

**Supplementary Fig. S2:**
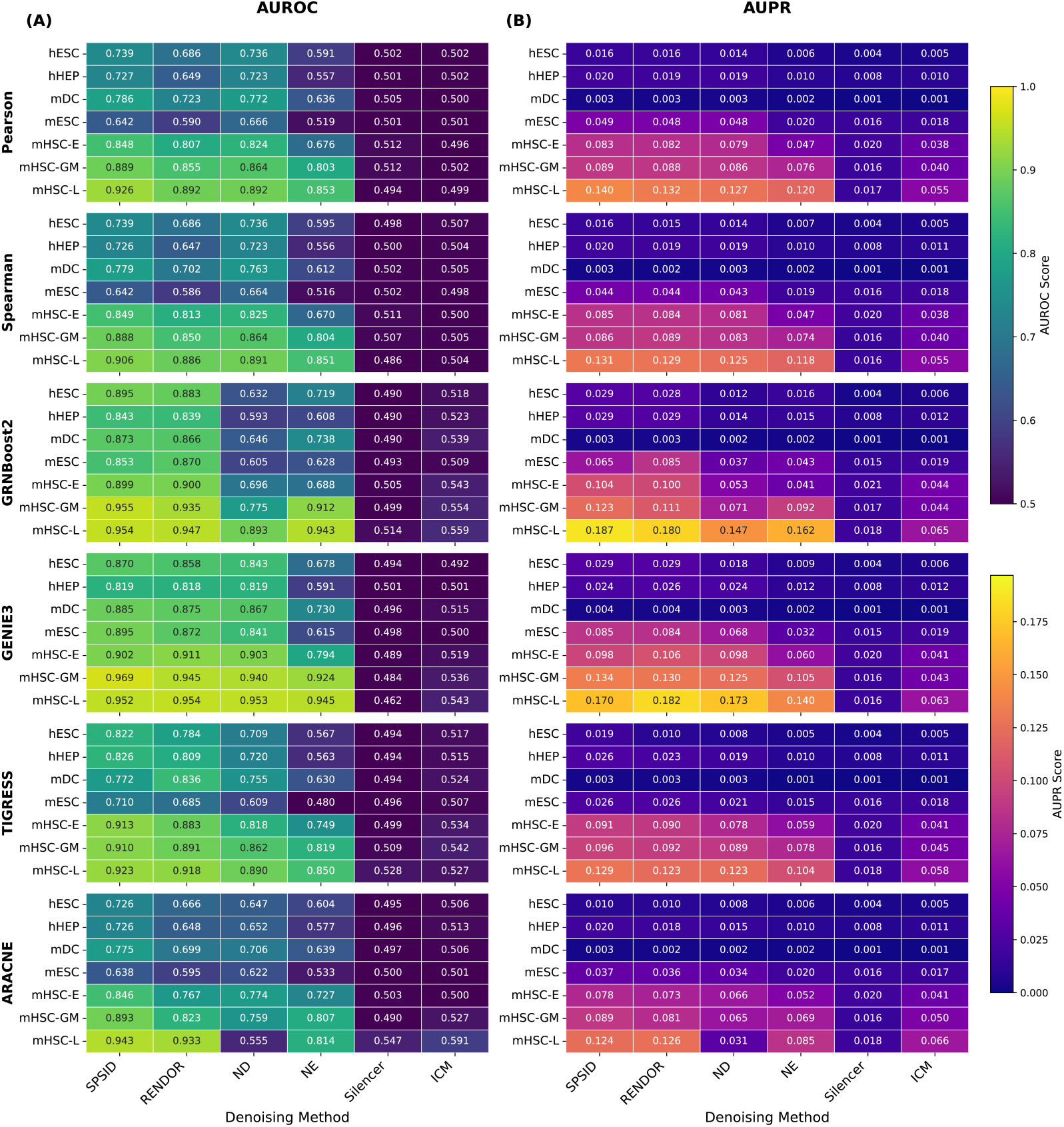
scRNA-seq benchmark performance. Facet heatmap displaying AUROC (A) and AUPR (B) scores. Each row-group represents one of the 6 upstream inference methods, and each row within a group represents one of the 7 scRNA-seq datasets. Columns show the SPSID and 5 other denoising methods.

**Supplementary Fig. S3:**
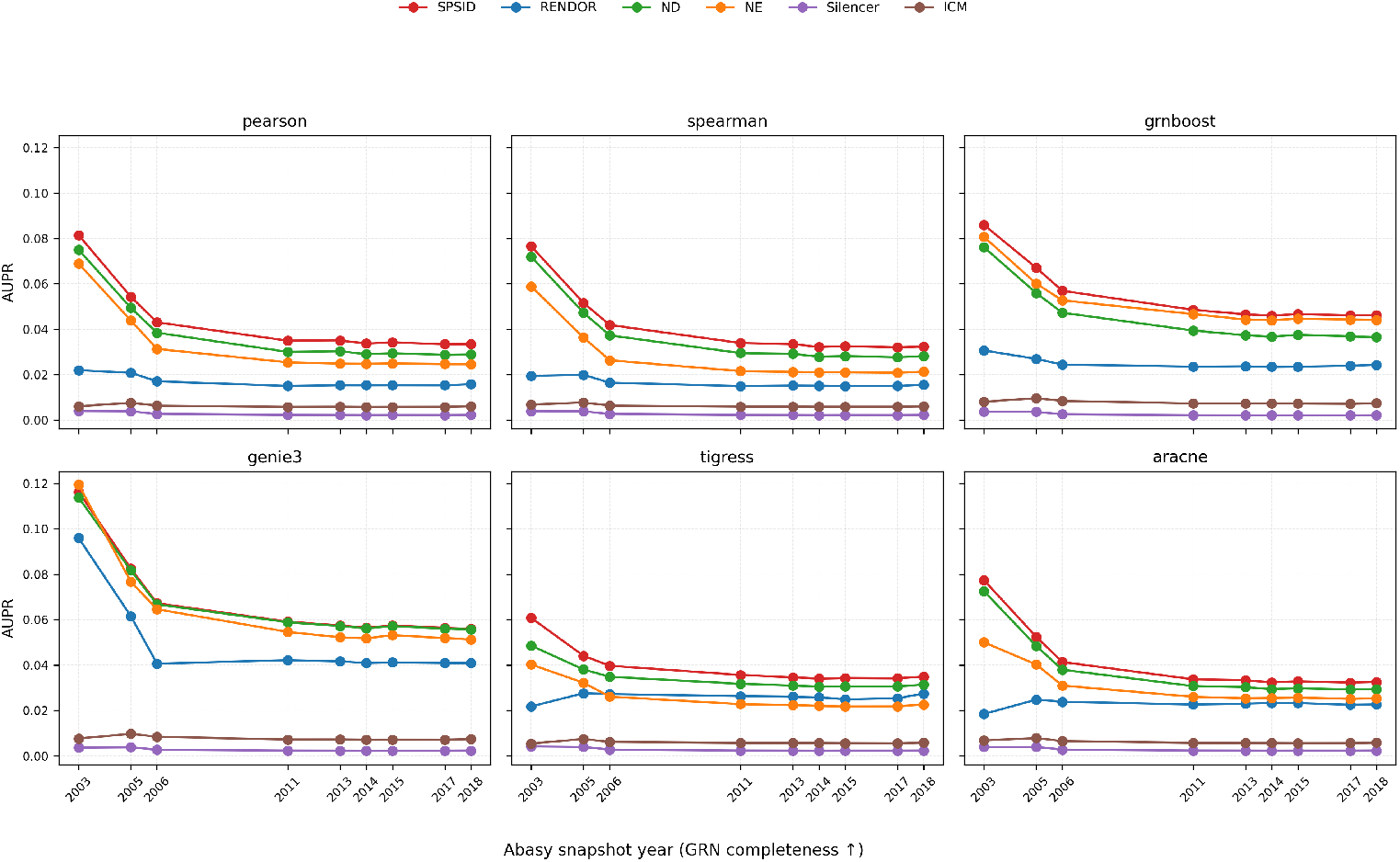
AUPR of denoising methods as a function of Abasy *E. coli* snapshot year (used as a proxy for GRN completeness). Rows correspond to upstream inference methods (Pearson, Spearman, GRNBoost2, GENIE3, TIGRESS, ARACNE); each panel compares SPSID with five competing denoisers. All methods are evaluated on the same DREAM5 Network 3 expression data and candidate edge set, but against Abasy-based gold standards of increasing completeness (2003–2018).

**Supplementary Fig. S4:**
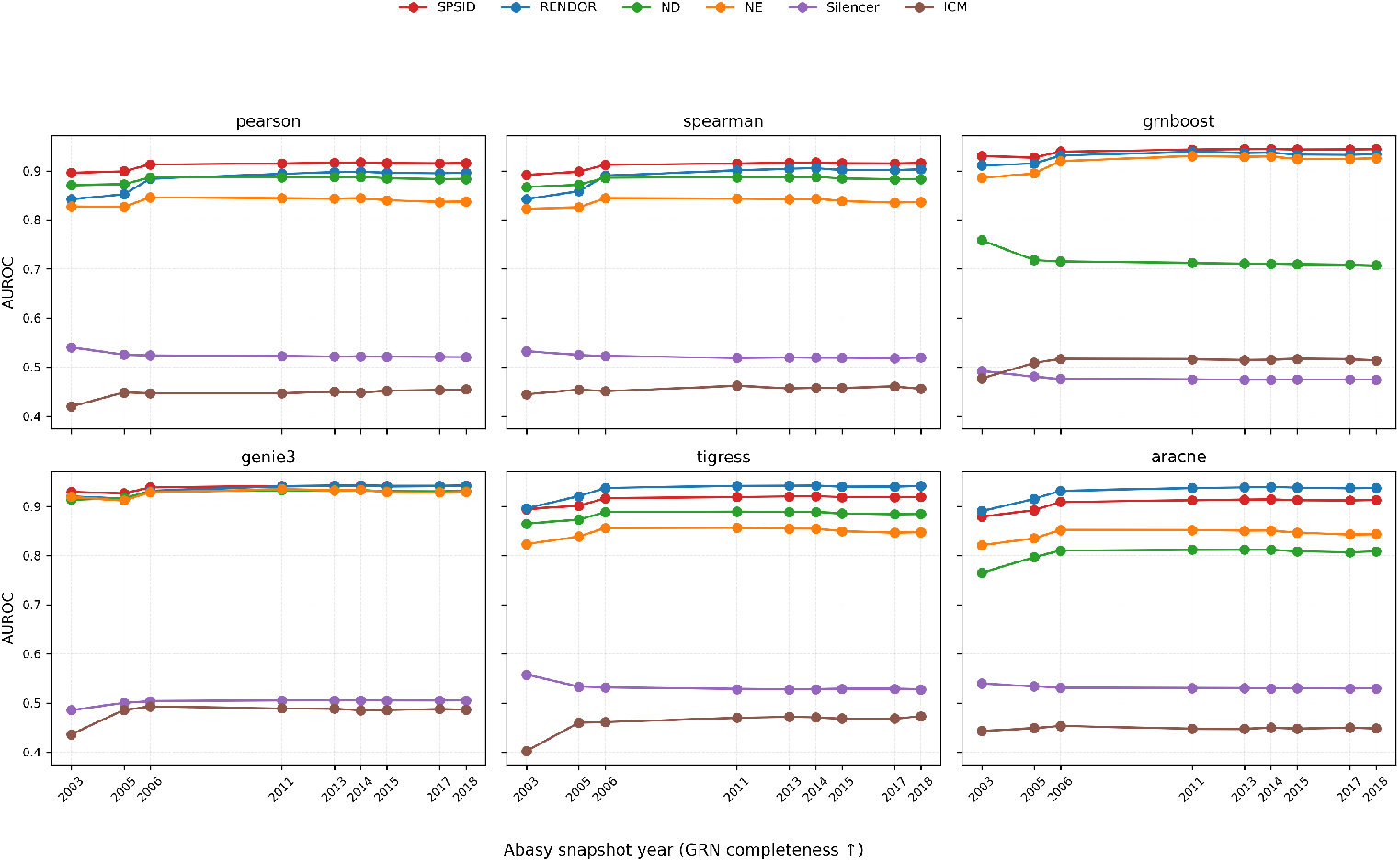
AUROC of denoising methods as a function of Abasy *E. coli* snapshot year. AUROC values are much more stable across snapshots than AUPR, and SPSID consistently matches or outperforms the best competing denoiser for all upstream backbones and completeness levels.

## Notes

### Competing Interest Statement

The authors have declared no competing interest.

